# A critical role for VE-cadherin in regulating actin dynamics during endothelial maturation and non-inflammatory activation via a tension-sensitive intermediate state

**DOI:** 10.1101/2025.08.05.668624

**Authors:** Jonas Franz, Maria Odenthal-Schnittler, Jan Philip Kipcke, Jochen Seebach, Zahra Labbaf, Franziska Merten, Vesna Bojovic, Johannes Eble, Erez Raz, Milos Galic, Hans Schnittler

## Abstract

Epithelial and endothelial monolayers maintain homeostasis by adapting to physiological stimuli and injury through conversion processes that remain incompletely understood. Using endothelial cell cultures (HUVEC), we investigate how monolayer maturation and non-inflammatory remodeling are molecularly regulated. Maturation involves reduced cell perimeter causing increased junctional VE-cadherin, which recruits junctional actin and integrins, establishing a quiescent, stable monolayer. Remarkably, we identify a previously unrecognized, rapid and reversible *intermediate-state*, marked by VE-cadherin linearization and actomyosin relaxation via MLC-dephosphorylation, that emerges during non-inflammatory activation triggered by onset or increase in shear stress. This novel *intermediate-state* enhances junctional actin and integrin recruitment, strengthening barrier-function while protecting endothelial cells from overstimulation and mechanical damage. Re-phosphorylation of MLC dissolves junctional actin and induces formation of *junction-associated-intermittent-lamellipodia* (JAIL), enabling cell shape change and arterial phenotype conversion. Overall, loss of actomyosin tension and junctional VE-cadherin-concentration defines actin recruitment and reveals a tension-sensitive, cell-protective intermediate state that primes endothelial remodeling, offering an expanded model for mechano-transduction and shear stress adaptation.

## Introduction

The internal and external surfaces of the human body, including the skin, intestinal mucosa, and cardiovascular and lymphatic systems, are lined by specialized layers of epithelial or endothelial cells that maintain homeostasis. These cell types exhibit distinct morphological and functional specializations, guided by complex genetic and epigenetic programs ^1–4^. For example, epithelial layers act as physical barriers that protect the body’s surface, facilitate nutrient absorption, and defend against pathogens. In contrast, endothelial monolayers line blood and lymphatic vessels, regulating perfusion and fluid exchange, functions essential for processes such as angiogenesis, wound healing, and inflammation. Despite these differences, both cell layers share key features such as apico-basal polarity, anchorage-dependent growth, and intercellular adhesion complexes that establish and maintain barrier function and are involved in regulating cell and junction remodeling. These structural elements work together to preserve tissue architecture and function, even during continuous turnover and regeneration ^5, 6^. Importantly, they also display adaptability to changing physiological demands ^7–13^. For instance, in response to fluid shear stress, endothelial cells elongate and align to acquire an arterial phenotype, a process that requires non-inflammatory activation ^14, 15^.

Tissue homeostasis is frequently challenged by inflammation or physical, chemical, or metabolic injury. Surface repair can result in three possible outcomes: complete restoration (restitutio ad integrum), scar formation known as defect regeneration, or chronic inflammation, where impaired healing leads to persistent tissue dysfunction and long-term diseases such as atherosclerosis ^16–19^. Understanding the transitions between resting and active states within these cell layers is therefore crucial to elucidate normal healing and to identify cellular targets for chronic disease intervention. Underlying these transitions are remodeling processes controlled by mechanical forces, growth factors, and inflammatory mediators. These factors regulate cell morphology, motility, and proliferation, processes that are mainly driven by actin cytoskeleton dynamics, intercellular junctions, and cell-matrix adhesions ^20–33^. These cellular mechanisms can produce either a pro-inflammatory, active phenotype during inflammation^34,35^ or a non-inflammatory, adaptive phenotype e.g. during physiological responses such as shear stress. However, the molecular coordination of these transitions, particularly between quiescent and active monolayer states, is still not fully understood.

To address this knowledge gap, we employ human umbilical vein endothelial cells (HUVECs) as a model system. HUVECs readily form confluent monolayers, undergo maturation, and respond to fluid shear stress with cell elongation and alignment. Their human origin, lack of ethical concerns, broad availability, and suitability for genetic manipulation and high-resolution live-cell imaging ^36^ make them ideal for studying transitional states in endothelial biology.

A central aspect of monolayer remodeling involves actin cytoskeleton dynamics and its interactions with adhesion complexes. In endothelial cells, actin polymerization generates branched networks that form membrane protrusions, as well as linear, myosin-containing bundles such as stress fibers and junctional actin ^37–40^. At intercellular junctions, actin interacts with VE-cadherin and its associated proteins to regulate adhesion and drive membrane protrusion formation, mechanisms that are essential for maintaining endothelial integrity and enabling remodeling ^21, 41, 42,23, 43,44,45–47^. With respect to membrane protrusions, junction-associated intermittent lamellipodia (JAIL) form at sites of low VE-cadherin concentration, such as intercellular gaps, directly creating new VE-cadherin adhesion sites. This dynamic interplay allows intercellular junctions to be remodeled while the overall monolayer maintains its basal integrity ^9, 22, 48^ ^35^. Furthermore, β1 and αvβ3 integrins anchor cells to the extracellular matrix, and their function is modulated by both cytoskeletal organization and VE-cadherin-mediated adhesion^20, 49, 50^.

In this study, we investigate two key processes that regulate the dynamics of the endothelial monolayer: maturation during cell growth and activation in response to shear stress. Our results identify VE-cadherin concentration at junctions, together with actomyosin-mediated tension, as central drivers of actin and integrin reorganization. Notably, we describe for the first time a novel tension-sensitive transient intermediate state that arises during the non-inflammatory conversion to an arterial phenotype. This state stabilizes endothelial junctions through VE-cadherin-dependent recruitment of actin and integrins, shielding cells from the sudden onset or increase of shear stress and priming them for subsequent remodeling. Together, these components, and the intermediate state, integrate mechanical and molecular signals to coordinate the balance between endothelial stability and plasticity, fundamental principles of cell biology.

## Results

### Junctional VE-cadherin shapes actin dynamics and directs integrin localization during the maturation of human umbilical vein endothelial cell monolayers

In order to investigate the maturation process of endothelial monolayers, we analyzed the dynamics of human umbilical vein endothelial cells (HUVECs) during growth, employing phase-contrast live-cell imaging and high-resolution structured illumination microscopy (SIM) over a 120-hour culture period. At an initial seeding density of 10^4^ cells/cm^2^, cells adhered to the substrate predominantly as single cells or small aggregates within two hours of plating (see Supplementary Fig. S1a). During this early phase, cells exhibited a low mean migration velocity of approximately 0.2 µm/min, showing limited mean Euclidean and accumulated migration distances (Figures 1a, 1b). Notably, individual cells demonstrated greater migratory activity than small cell clusters at this stage (HS, personal communication).

**Figure 1.**
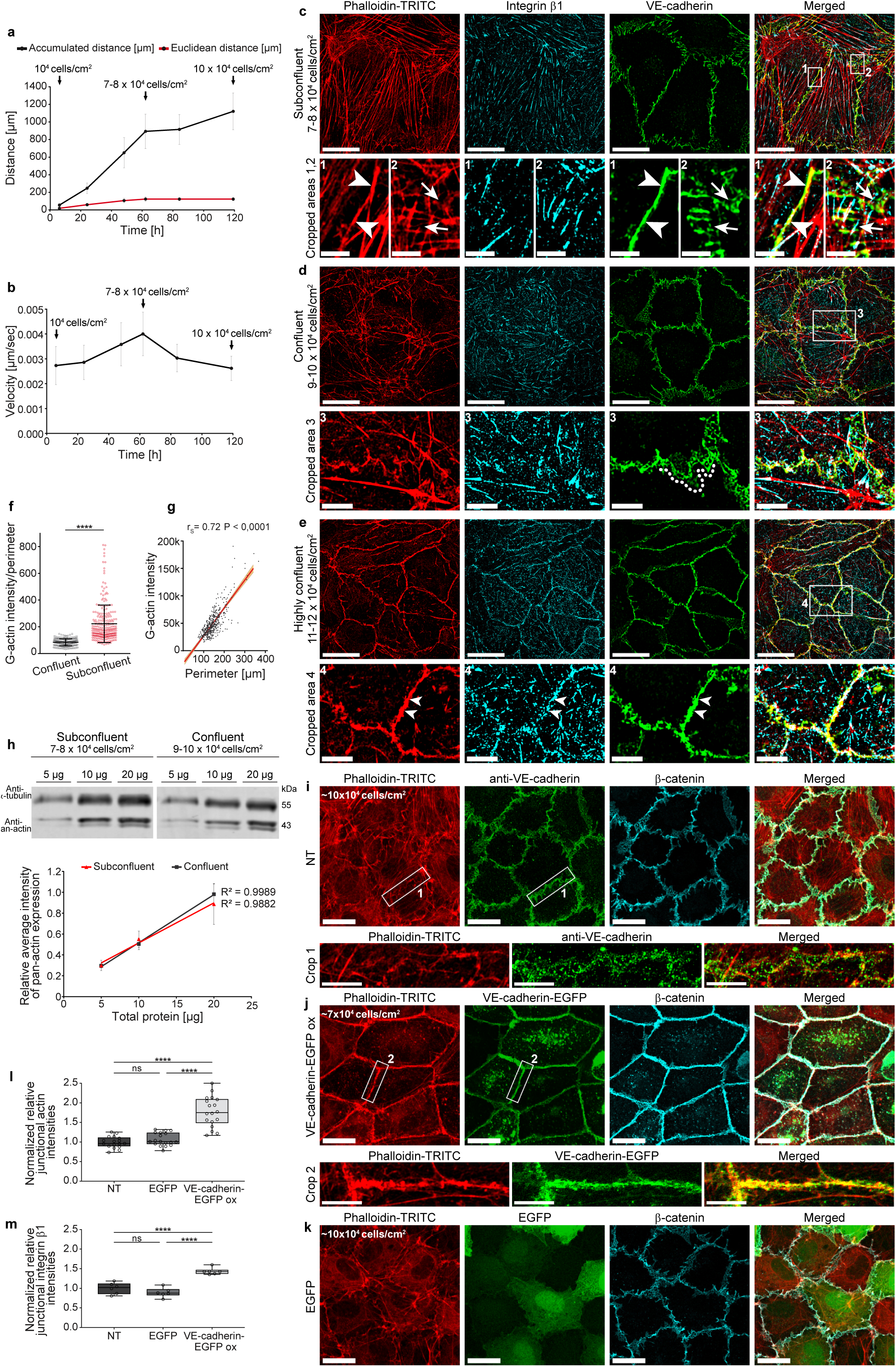
VE-cadherin, actin and integrin patterns predict HUVEC dynamics. **(a,b)** Cell-density-dependent migration. **(a)** Accumulated distance, Euclidean distance; **(b)** velocity. N = 150 cells, 3 independent experiments. Note: cell migration velocity temporarily increases with increasing cell density. **(c-e)** Density-dependent distribution of actin filaments, VE-cadherin, and integrin β1 in HUVECs visualized by structured illumination microscopy (SIM) and the one-drop assay. **(c, d)** At a density between 7-8 x 10^4^ and 9-10 x 10^4^ cells/cm^2^, the heterogeneous distribution of VE-cadherin includes interrupted junctions (Crop 2, arrows), linear junctions (Crop 1, arrowheads), and plaques indicating JAIL (Crop 3, dotted line). Integrin β1 is localized overall at focal contacts. Note: actin filaments partially colocalize with both VE-cadherin and integrin β1. (**e**) Further culturing of 9-10 x 10^4^ cells/cm^2^ for 8 days increased the cell density to 11-12 x 10^4^. This promoted junction maturation, characterized by a linear (condensed) VE-cadherin distribution overlapping junctional actin and recruitment of integrin β1 to cell-cell contact areas (Crop 4, arrowheads). Note: stress fibers were largely absent. Overviews: scale bar, 20 µm; Crops 1, 2: 3 µm; Crops 3, 4: 5 µm. (**f, g**) The ratio of G-actin to cell perimeter shows a clear density dependency, analyzed by DNase I immunolabeling using the one-drop assay. Cell perimeters were measured with the Cell Border Tracker (CBT). (**f**) G-actin/perimeter ratio in sub-confluent (n = 244 cells) and confluent (n = 208 cells) cultures; Mann-Whitney test. (**g**) Correlation between DNase I labeling and cell perimeter (n = 486 cells; rₛ = Spearman correlation coefficient). (**h**) Density-dependent analysis of total actin by Western blot in sub-confluent and confluent cells as indicated. Different amounts of total cellular protein (5, 10, and 20μg) were probed with an anti-pan-actin antibody, followed by densitometry and determination of r^2^. α-Tubulin served as a loading control (n= 4 independent experiments). (**i-m**) Effect of VE-cadherin overexpression on actin filament distribution analyzed by quantitative immunolabeling with LSM and SIM. (**i**) Control cells at 10 x 10^4^ cells/cm^2^ and (**j**) VE-cadherin-overexpressing cells. Note: complete loss of stress fibers and strong junctional recruitment of actin in VE-cadherin-overexpressing cells, while (**k**) overexpression of EGFP alone, even at higher density (10 x 10^4^ cells/cm^2^), maintained the heterogeneous pattern (overview scale bar, 20 µm; Crop 1, SIM: 5 µm). (**l**) Quantification of junctional actin and (**m**) junctional integrin β1 in VE-cadherin-EGFP-overexpressing cells and controls as indicated using the CBT (one dot = 15 cells; one-way ANOVA). See integrin distribution in Supplementary Figure 1d. All results shown are from at least 3 independent experiments. Error bars indicate ± SEM (ns ≥ 0.05, **** p < 0.0001).

Over the subsequent 60 hours, cell density increased markedly to reach 7-8 × 10^4^ cells/cm^2^. This was accompanied by a significant increase in migration velocity and distance travelled, resulting in the formation of a complete monolayer (Figures 1a, 1b, Supplementary Fig. S1a). However, the junctions remained immature which was confirmed by SIM following immunolabelling of actin, integrins and VE-cadherin using the one-drop culture assay - an approach that enables the analysis of molecular patterns across varying cell densities under uniform culture conditions (Figure 1c, Supplementary Fig. S1b). As the cell density further increased up to 9-10 x10^4^ cells/cm^2^, a reduction in stress fibers was observed, although a few were still present and terminated at integrin β1-positive structures throughout the cells (Figure 1d). Additionally, an irregular VE-cadherin distribution was evident, indicating ongoing dynamic junctional remodelling (Figure 1d). This is consistent with the observed increases in migration velocity and accumulated distance travelled (Figures 1a, 1b). However, the Euclidean distance increased only slightly (Figure 1a), suggesting that cell migration remained largely random rather than directionally polarised.

As the cell density increased, reaching a maximum of 11-12 x 10^4^ cells/cm^2^ over the 120-hour culture period, the monolayer acquired a characteristic cobblestone morphology (Supplementary Fig. S1a). At this stage, both cell migration and junctional dynamics decreased to low levels (Figure 1a, 1b). Consistently, SIM analysis revealed features of cell and junction maturation, which included the depletion of stress fibers and the moderate development of junctional actin, with integrin β-1 being recruited to these sites (Figure 1e).

To better understand actin dynamics during monolayer maturation, we examined G-actin levels at various cell densities. G-actin provides the monomers necessary for the polymerization of actin filaments, which is an essential process for cellular protrusions, migration and junction remodeling ^30, 37, 51^. Using the DNase I assay (Supplementary Fig. S1b), we found that G-actin levels decreased as cell density increased, resulting in reduced cell migration and circumference (Figure 1f, 1g, Supplementary Fig. S1b). However, total actin expression remained unchanged (Figure 1h), which is consistent with *in vivo* findings ^48^. The reduction in G-actin coincided with the formation of junctional actin structures and integrin β1 recruitment to cell junctions (Figure 1e; Supplementary Fig. S1b). Overall, these results demonstrate that migratory behavior shifts in response to changes in cell density, driven by the reorganization of the actin cytoskeleton in coordination with integrin- and VE-cadherin-based adhesion complexes.

### VE-cadherin overexpression forces a mature monolayer phenotype, even in sub-confluent cells

As cell density increases, cell circumference decreases, but total VE-cadherin expression remains unchanged ^35,52^. This leads to a higher concentration of VE-cadherin at cell junctions. We hypothesized that this elevated junctional VE-cadherin level would facilitate junctional actin assembly and stabilize both interendothelial and cell-substrate adhesions. To test this hypothesis, VE-cadherin-EGFP was overexpressed by about 2-3 times ^35^ in sub-confluent HUVECs (7 x 10^4^ cells/cm^2^). In control cells, stress fibers and diffuse integrin distribution were observed alongside a discontinuous VE-cadherin pattern (Figures 1i, Supplementary Figs. S1c). In contrast, VE-cadherin-overexpressing cells formed a stable VE-cadherin/catenin complex, as demonstrated by colocalization with β-catenin. Furthermore, this resulted in the development of a pronounced linear junctional actin (Figure 1j,1l), as well as a marked increase in integrin β1 at this site (Figure 1m, Supplementary Fig. S1c, S1d). Importantly we found a total loss of cytoplasmic stress fibers and focal adhesions. Overexpression of EGFP alone did not induce these changes (Figures 1k, 1I, 1m). These results demonstrate that increased junctional VE-cadherin recruits actin to cell-cell contacts and reorganizes the actin cytoskeleton, producing an endothelial phenotype similar to mature venous and arterial endothelium ^18, 53–56^. These findings highlight the pivotal role of VE-cadherin in regulating endothelial maturation and quiescence through the modulation of actin dynamics and integrin distribution.

### Shear stress-induced transition to an arterial phenotype occurs through an intermediate cellular state

Next, we investigated the dynamics and mechanisms underlying the conversion of endothelial cells from a quiescent to an active state, forced by fluid shear stress. To this end, we applied unidirectional fluid shear stress (U-FSS) to endothelial monolayers using a the cone-and-plate BioTechFlow system (BTF system), (Figure 2a), which enables phase-contrast and fluorescence live-cell imaging simultaneous with barrier function measurements, followed immunolabelling and biochemical assays ^57–59^.

**Figure 2.**
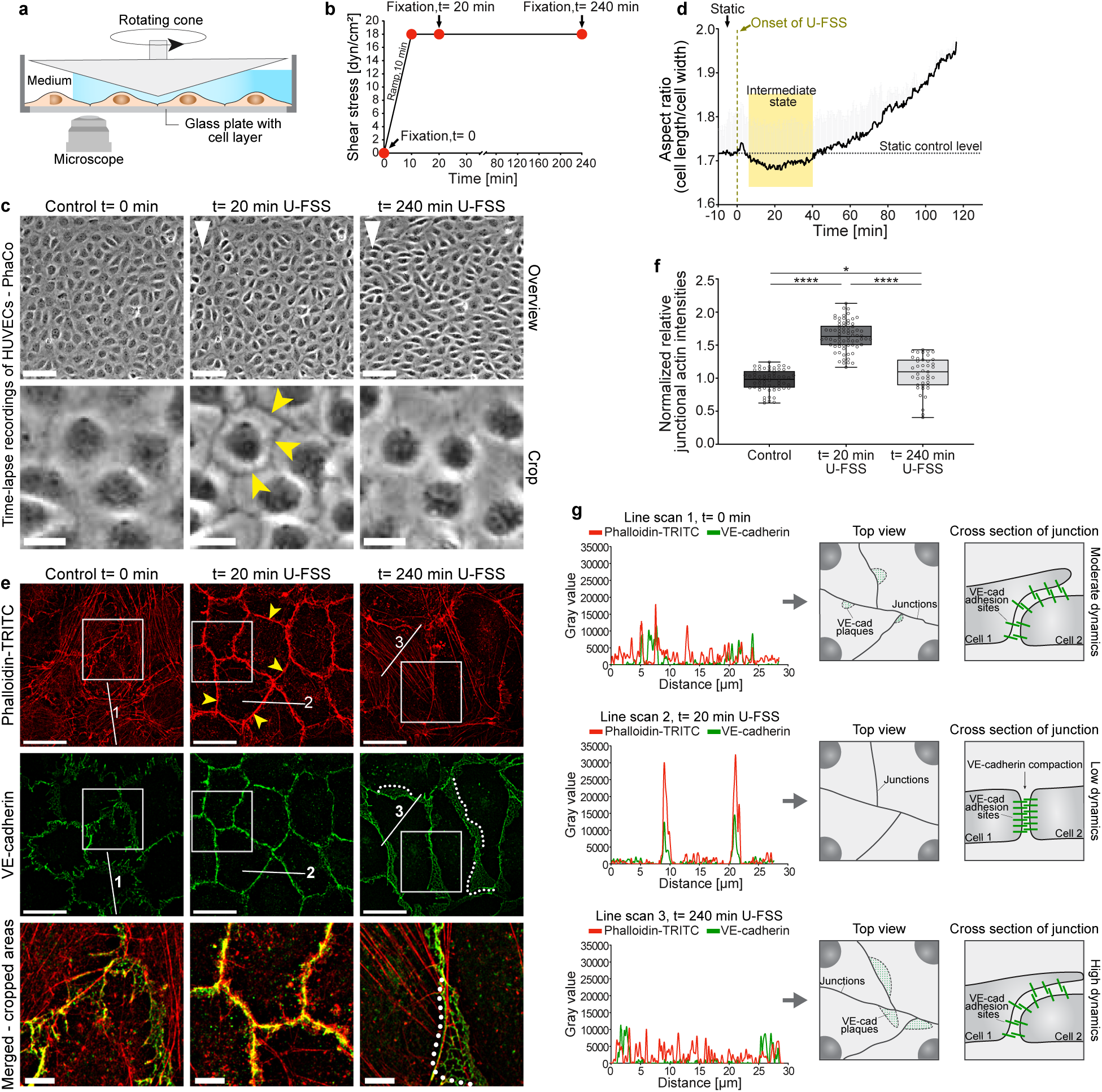
Shear stress drives EC remodeling through an intermediate state. HUVECs (9-10 x 10^4^ cells/cm^2^) were exposed to 18 dyn/cm^2^ of unidirectional fluid shear stress (U-FSS) for various time points and then immunolabelled as indicated. (**a**) Scheme of the cone-plate rheological BTF in vitro system. (**b**) Shear stress profile used in the experiments; red dots mark the time points shown below. (**c**) Phase-contrast (PhaCo) time-lapse images. Note the increased border contrast (yellow arrowheads) within 20 minutes under U-FSS, followed by cell shape changes and loss of junction contrast (overview scale bar: 100 µm; crops: 20 µm; see Supplementary Movie 1). White arrowheads indicate the U-FSS direction. (**d**) Aspect ratio (AR) changes during U-FSS, quantified by PhaCo microscopy and a custom software tool. U-FSS caused a transient cobblestone-like shape with decreased AR within 20 minutes, followed by shear-induced cell elongation indicated by increased AR (n ≈ 4000 cells/time point). (**e**) Structured illumination microscopy (SIM) images of HUVECs exposed to U-FSS at t= 0 min, t= 20 min and t= 240 min, labelled for VE-cadherin and actin filaments using phalloidin-TRITC (overview scale bar: 20 µm; cropped areas: 5 µm). Note the prominent recruitment of junctional actin (arrowheads) and the loss of stress fibers after 20 minutes of U-FSS. Prolonged U-FSS exposure resulted in increased junctional JAIL dynamics, indicated by VE-cadherin plaque formation (dotted line). Note, the loss of junctional actin, interrupted VE-cadherin and the reformation of stress fibers. (**f**) Quantification of relative junctional actin intensities prior to U-FSS (control; n ≈ 900 cells), after 20 min (n ≈ 1080 cells) and 240 min of U-FSS (n ≈ 615 cells) using the CBT (n ≈ 15 cells/dot; one-way ANOVA). (**g**) Line plots of junctional actin and VE-cadherin distribution as shown in (e). The schematic top views and cross-sections illustrate the proposed underlying junctional dynamics. (**d,f**) Error bars represent ± SEM (*p < 0.05, ****p < 0.0001). Results of three independent experiments are shown.

Control monolayer of HUVEC at a density of 10 x 10^4^ cells/cm^2^ displayed heterogeneous VE-cadherin pattern and moderate junctional actin, with few stress fibers. U-FSS was gradually increased to 18 dyn/cm^2^ over 10 minutes to mimic physiological adaptation to increased blood flow (Figure 2b, Supplementary Movie 1). Phase-contrast live-cell imaging revealed that within 20 minutes, cell borders became more distinctly visible due to increased contrast, accompanied by the formation of a cobblestone-like cell pattern (Figures 2c, 2d, Supplementary Movie 1). This morphological change coincided with VE-cadherin reorganizing into homogeneous linear, condensed pattern. Impressively, junctional actin became prominent while stress fibers were completely absent (Figures 2e, 2f, 2g, Supplementary Fig. 2a).

Due to its transient nature, we classified this phenotype as an intermediate state. However, with prolonged U-FSS exposure, the actin ring gradually disassembled, and stress fibers reformed (Figures 2c, 2e, 2f). VE-cadherin dynamics increased through actin-mediated junction-associated intermittent lamellipodia (JAIL) formation, as evidenced by the appearance of VE-cadherin plaques (Figures 2e and 2g). This process ultimately led to cell elongation and alignment in the direction of shear stress, as frequently shown ^58, 60–62^. Notably, the loss of junctional actin observed under sustained fluid shear stress (FSS) also occurs when U-FSS is stopped after 20 min (HS personal communication). This dramatic and transient junctional actin remodeling prompted us to evaluate the interplay between the recruitment of actin and the distribution of integrins, as both are key molecules involved in cell-substrate adhesion and mechano-transduction^50^. Indeed, integrins β1 and avβ3 strongly colocalized with recruited junctional actin, while cytoplasmatic integrin-positive focal adhesions were completely absent, compared to controls (Figures 3a, 3b, 3d, 3e, Supplementaryw Fig. 2b), whereas overall integrin β1 expression remained unchanged (Figure 3f). Additionally, TIRF microscopy confirmed colocalization of junctional actin and integrin αvβ3 at the basal cell side (Supplementary Fig. 2c). These data suggest that the localization of integrins depends on actin filaments. Consistent with this, prolonged U-FSS results in changes to integrin distribution that occur alongside junctional actin disassembly and stress fiber formation (Figure 3a). This is accompanied by shear stress-induced endothelial remodeling through actin-driven JAIL. JAIL typically overlap with adjacent cells and directly form new VE-cadherin adhesion sites ^21^ thereby enabling dynamic junction remodeling while preserving a certain degree of monolayer integrity. In addition to this frequently observed mechanism, we found using SIM and TIRF microscopy that, in some cases, integrins β1 and αvβ3 are prominently localized at the protrusive front of JAIL structures during their recruitment after 20 minutes (Figure 3b). This suggests that JAIL might extend beneath adjacent cells. Following the disassembly of junctional actin and the formation of enhanced JAIL structures that drive cell shape changes, integrins were notably absent from these structures (Figure 3c). Instead, they accumulated predominantly at the ends of stress fibers, which is consistent with previous reports ^63, 64^.

**Figure 3.**
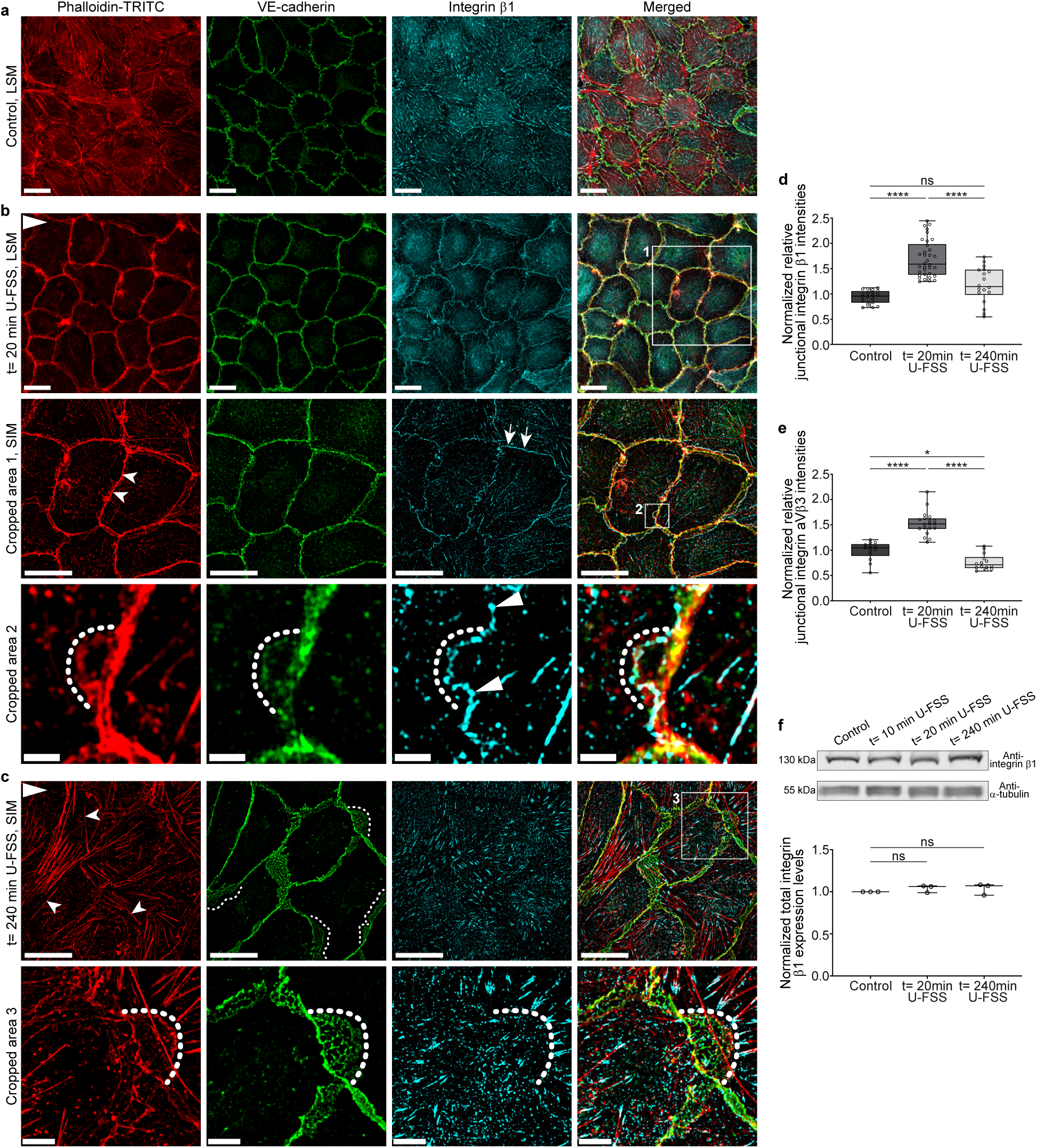
Shear stress-induced transient junctional actin recruitment was accompanied by integrin β1 and integrin α_V_β3 localization at cell junctions. (**a-c**) LSM and SIM of confluent HUVECs (9-10 x 10^4^ cells/cm^2^) exposed to 18 dyn/cm^2^ U-FSS for (**a**) t= 0 min, (**b**) t= 20 min, and (**c**) t= 240 min, stained with phalloidin-TRITC, VE-cadherin, and integrin-β1. Scale bars = 20 µm. The U-FSS direction is indicated in the upper left corner of each figure. (**b**) Prominent junctional actin formation (arrowheads) accompanied by integrin β1 recruitment (arrows) and corresponding VE-cadherin linearization after 20 min. Higher magnification by SIM (Crop 1, scale bar 20 µm). Note: small JAILs with remodeling actin-driven VE-cadherin are positive for integrin β1 at the extending JAIL borders (Crop 2, SIM, scale bar 2 µm). (**c**) Sustained U-FSS led to reappearance of stress fibers (arrowheads) connected to integrin β1-positive focal adhesions, while large JAILs, indicated by VE-cadherin plaques (Crop 3, dotted lines), formed and drove cell elongation. Time-dependent junctional recruitment of (**d**) integrin β1 and (**e**) integrin α_V_β3 was quantified using the CBT in immunolabeled HUVECs as indicated. See Supplementary Figures 2b and 2c for integrin α_V_β3 labelling. One dot= 15 cells; one-way ANOVA. (**f**) Representative Western blot and quantification of integrin β1 from confluent HUVECs exposed to 18 dyn/cm^2^ U-FSS. α-Tubulin served as the internal loading control; one-way ANOVA. Data are from 3 independent experiments. Error bars indicate ± SEM (ns ≥ 0.05, * p < 0.05, **** p < 0.0001).

### A temporal blueprint of endothelial junction adaptation to shear stress revealed by live-cell fluorescence microscopy

To better resolve the temporal dynamics of protein remodeling under unidirectional fluid shear stress (U-FSS), we moderately expressed LifeAct-mCherry and VE-cadherin-EGFP in HUVEC cultures seeded at 10 x 10^4^ cells/cm^2^ (Supplementary Figs. 3a-3c). Time-lapse recordings revealed that stress fibers began to dissociate after just 8-10 minutes of U-FSS application, while a prominent junctional actin ring was formed within 20 minutes (Figure 4a, Supplementary Movie 1, middle panels). Accordingly, the VE-cadherin-EGFP pattern dynamically shifted from a broad, heterogeneous to a stable linear arrangement within 20 minutes (Figure 4c, Supplementary Movie 1, right panels). Consistently, quantitative analysis of VE-cadherin dynamic displacement using a cell border tracker revealed a 20% decrease during the proposed intermediate cellular state (Figures 4d, 4e). This phenotype closely resembles highly confluent, quiescent endothelial cells *in vitro* and large veins *in vivo* ^54, 55^, suggesting a transient, protective intermediate state in response to mechanical stress. A similar intermediate endothelial state was recently described following TNF-α treatment ^35^. We propose that the shear stress-induced intermediate state likewise protects cells from damage and primes them for remodeling.

**Figure 4.**
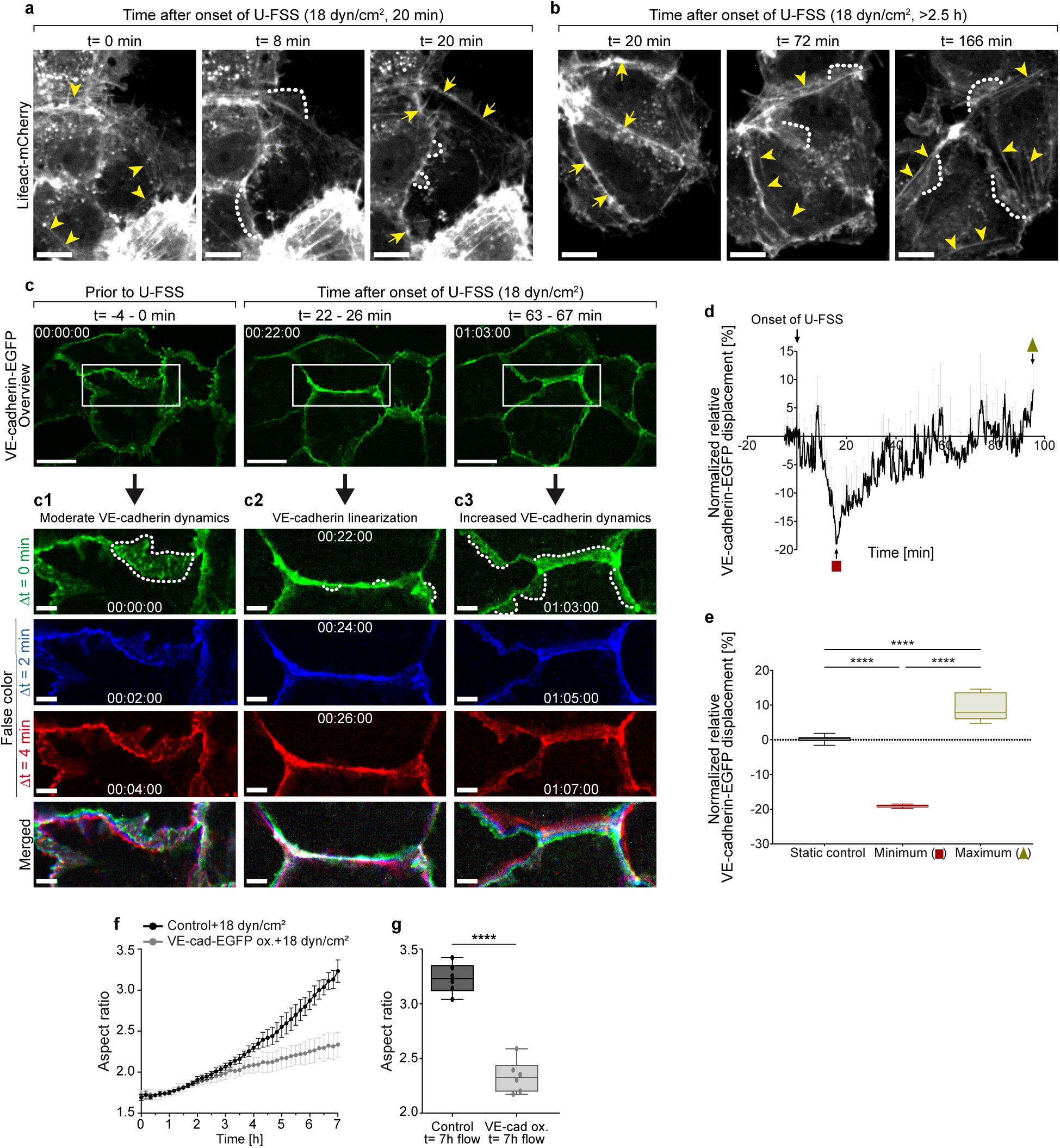
Actin and VE-cadherin dynamics under shear stress analyzed by live-cell imaging. (**a,b**) Time-lapse recordings of Lifeact-mCherry-expressing HUVECs (9-10 x 10^4^ cells/cm^2^) subjected to U-FSS at the indicated time points (scale bar: 10 µm). (**a**) Under static conditions (t=0), various actin filaments, including stress fibers (arrowheads), are visible. Upon U-FSS onset, stress fibers dissociate while a prominent transient junctional actin ring forms by t=20 min (arrows). (**b**) Continued U-FSS leads to the dissociation of junctional actin while stress fibers (arrowheads) reappear (compare Supplementary movie 1,2). (**c**) Time-lapse recordings of moderately VE-cadherin-EGFP-expressing confluent HUVECs exposed to U-FSS (scale bar: 20 µm). (**c1-c3**) Cropped areas (scale bar: 5 µm) show VE-cadherin-EGFP before U-FSS (**c1**), at 22-26 minutes (**c2**), and at 63-67 minutes (**c3)** after U-FSS, displayed in false colors for times Δt= 0 (green), Δt= 2 min (blue), and Δt= 4 min (red). (**c2**) Note the transient linearization of VE-cadherin-EGFP after 22-26 minutes of U-FSS, followed by a decrease in the size and frequency of JAIL (dotted lines). Prolonged U-FSS leads to (**c3**) increased VE-cadherin dynamics and enhanced JAIL formation (dotted lines) (time scale: hh:mm:ss); compare Supplementary Movie 1,2. (**d**) Determination of VE-cadherin-EGFP displacement (dynamics) within 100 min U-FSS based on CBT analyses. Note the transient decrease in dynamics after t≈ 20 min, followed by a continuous increase (n = 4 independent experiments). (**e**) Quantification of (d) comparing control (prior to U-FSS), minimum (t≈ 20 min) and maximum (t≈ 98 min); (n = 4 independent experiments; one-way ANOVA. (**f,g**) Aspect ratio analyses of VE-cadherin-EGFP overexpressing HUVECs and EGFP control cells subjected to 18 dyn/cm^2^ for 7 hours using phase-contrast microscopy and a custom analysis tool (n = 3 independent experiments; unpaired Student’s t-test). Error bars represent ± SEM (****p < 0.0001).

The ongoing shear stress reactivated cell junction dynamics, particularly the dissociation of junctional actin and the re-formation of stress fibers. Some of these stress fibers emerged directly from the junctional actin (Figure 4b, Supplementary Movie 2, left panel). This was accompanied by an increase in VE-cadherin displacement levels to above those of the control (Figures 4d, 4e), which subsequently drove cell elongation and alignment driven by increased JAIL formation (Figure 4c). Since JAIL formation requires local VE-cadherin dilution, we aimed to block this phenomenon by overexpressing VE-cadherin. Application of shear stress to these cells nearly completely inhibited shear stress-induced alignment (Figures 4f, 4g), demonstrating the significance of this mechanism for the shear stress-mediated adaptation of endothelial cells.

### Shear stress-induced actomyosin relaxation and tension loss triggers an intermediate state for endothelial remodeling

To elucidate the mechanisms underlying the transient intermediate state during endothelial remodeling induced by shear stress, we first examined the role of actomyosin dynamics. Cyclic contraction and relaxation, mediated by the phosphorylation of myosin light chain (MLC) of actin-myosin bundles, are well-established drivers of vascular remodeling ^65–68^. We confirmed that shear stress recruits the actin-myosin machinery by immunolabelling myosin IIa (Supplementary Fig. 4). Surprisingly, MLC dephosphorylation coincided with transient junctional actin accumulation, stress fiber reduction, and VE-cadherin linearization at cell borders (Figures 5a-5d), consistent with reduced JAIL formation (Supplementary movie 1, Figure 5a). Ongoing shear stress, however, reversed these effects, resulting in renewed MLC phosphorylation, dissociation of junctional actin, re-emergence of stress fibers and increased JAIL activity. This, in turn, promoted VE-cadherin dynamics and cell elongation (Figures 5a-5d, Supplementary movies 1 and 2).

**Figure 5.**
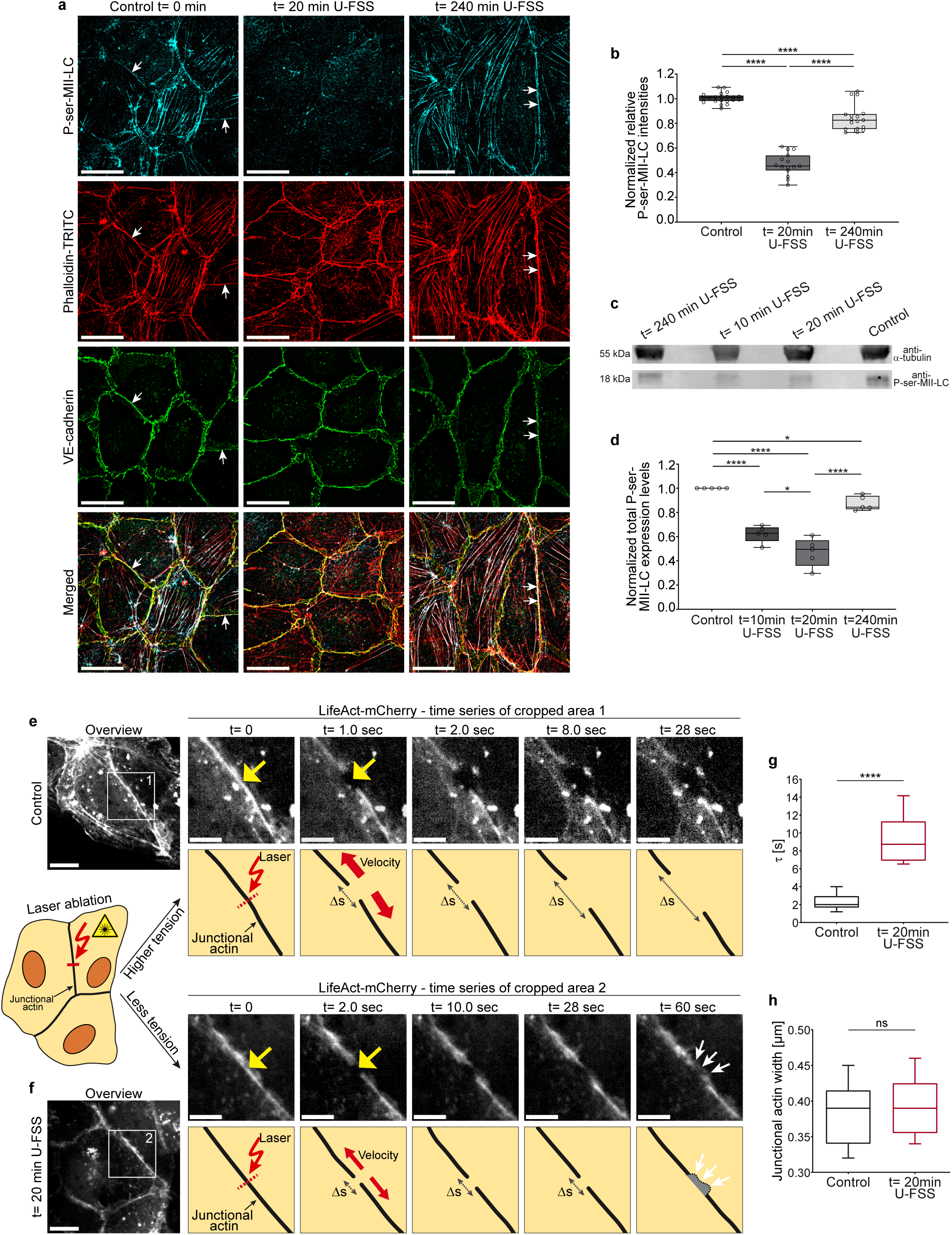
Shear stress triggers an intermediate state with junctional actin recruitment by transient myosin II light chain dephosphorylation-driven tension loss. (**a**) SIM images of phospho (ser19)-myosin II light chain (P-ser-MII-LC), phalloidin-TRITC and VE-cadherin -labelled confluent HUVECs (9-10 x 10^4^ cells/cm^2^) after application of 18 dyn/cm^2^ U-FSS for t= 0 min, t= 20 min and t= 240 min. Note, a transient myosin dephosphorylation is obvious after 20 min (scale bar: 20µm, U-FSS direction from the left). Arrows indicate MII-LC phosphorylation sites along junctional actin bundles in controls and during shape change at t= 240 min U-FSS. (**b**) Quantification of relative P-ser-MII-LC immunofluorescence intensities prior to U-FSS (control; n ≈ 315 cells), at t= 20 min (n ≈ 225 cells) and t= 240 min of U-FSS (n ≈ 255 cells); (n ≈ 15 cells/dot; one-way ANOVA). (**c,d)** Quantitative Western blot analysis of P-ser-MII-LC as indicated (one-way ANOVA). (**e,f**) Laser ablation of junctional actin in LifeAct-mCherry-expressing confluent HUVECs exposed to (**e**) static control conditions or (**f**) t= 20 min of U-FSS. Time-lapse recordings were performed of comparable junctional actin areas to follow the displacement behavior after ablation. Left, overviews (scale bar: 10 µm); right, time series of cropped areas 1 and 2 (note the different time points; scale bar: 5 µm) with matching scheme below (yellow arrows indicate the position of the laser spot; Δs: distance). (Area 2, t= 60 sec), white arrows indicate recovery of the incision site by actin protrusion (JAIL). (**g**) Quantification of laser ablation using the Kelvin-Body model to determine the elastic stiffness (τ) of the junctional actin in static control (n = 21 incisions) and at t= 20 min of U-FSS (n = 17 incisions). Less tension in junctional actin is indicated by increased τ levels after 20 min of U-FSS, while (**h**) the width of the analyzed junctional actin remains largely consistent; unpaired student’s t test. Data are from 3 independent experiments. Error bars indicate ± SEM (ns β 0.05, *p < 0.05, ****p < 0.0001).

To directly assess the importance of actomyosin-mediated tension, we performed laser ablation of junctional actin in HUVECs expressing LifeAct-mCherry that were subjected to either static conditions or 18 dyn/cm^2^ U-FSS for 20 minutes. Post-ablation time-lapse imaging revealed significantly less junctional actin displacement in shear-stressed cells than in static controls (Figures 5e, 5f). Kelvin-Voigt modelling revealed significant differences in the viscoelastic relaxation times (𝜏) of junctional actin (Figures 5g, 5h). Additionally, cutting the actin bundles after 20 minutes of shear stress resulted in the rapid formation of branched actin structures and JAIL under ongoing shear stress, indicating intact plasma membranes and active cytoskeletal remodeling (Figure 5f). These results are consistent with previous observations describing loss of membrane-cytoskeleton tension as a facilitator of cellular wound closure ^69^. In contrast, such a repair mechanism was rarely observed or occurred later in the static control (Figure 5e). These data confirm that shear stress induces the relaxation of junctional actin bundles, thereby highlighting the dynamic and reversible nature of the intermediate state in endothelial remodeling. To further analyze the impact of tension loss in actin bundles on VE-cadherin and actin distribution, we treated sub-confluent HUVECs (8 x 10^4^ cells/cm^2^) with the ROCK inhibitor Y27632. Within minutes, stress fibers dissociated, accompanied by an increase in JAIL formation (Figure 6a (2)). Over time, ROCK inhibition produced two distinct effects at junctional sites: in most cells, junctional actin became more prominent, while in others, JAIL formation persisted (Figures 6a (3), 6b; Supplementary Movie 3). In line with this effects immunofluorescence also revealed two distinct VE-cadherin patterns after ROCK application: densely organized VE-cadherin with strong junctional actin and integrin β1 recruitment, and irregular VE-cadherin with diffuse actin and integrin β1 (Figure 6c, for overview see Supplementary Fig.S5).

**Figure 6.**
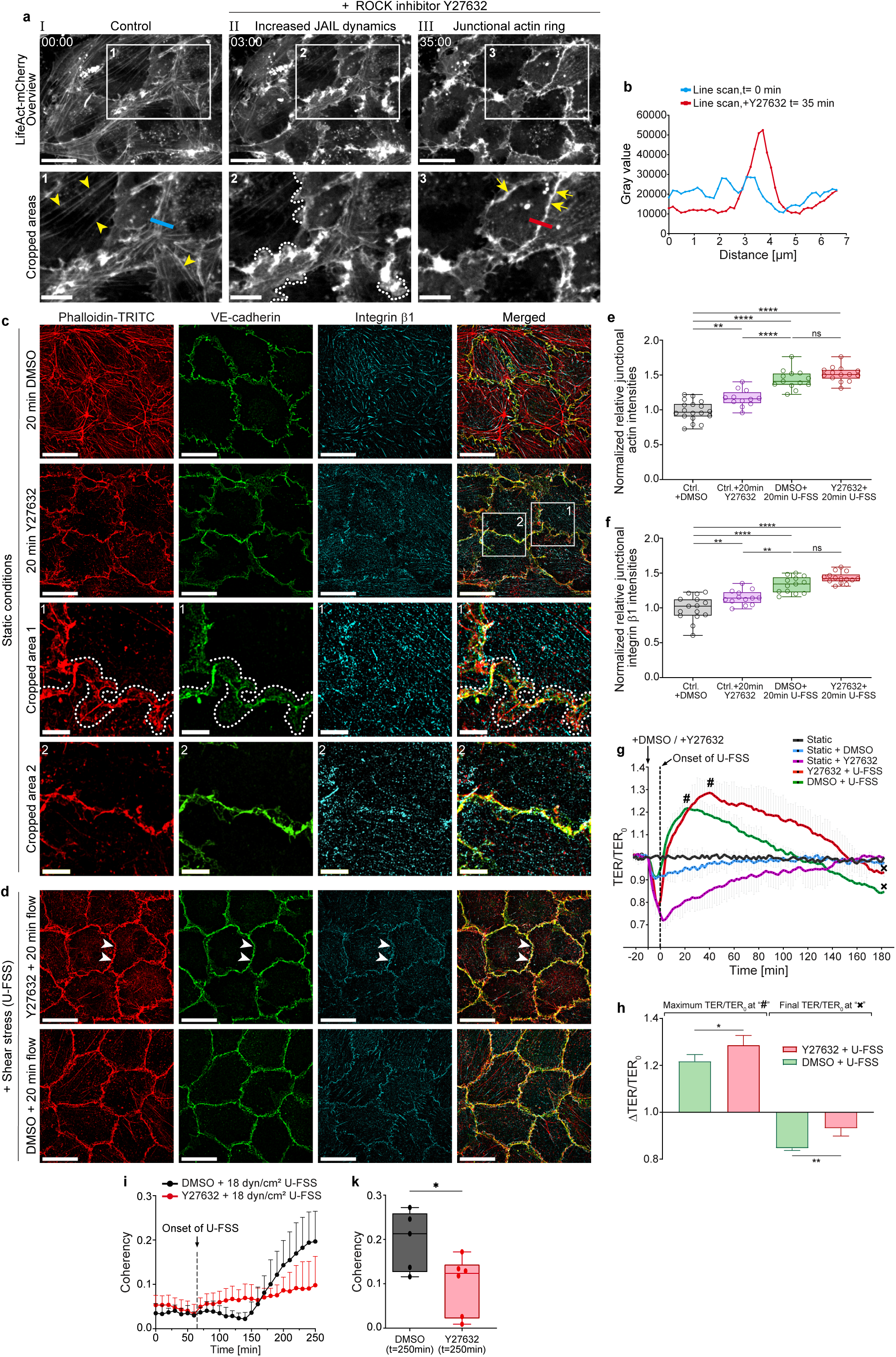
Differential effects of ROCK inhibitor-mediated selective relaxation and shear stress-induced cellular conversion. (a) Time-lapse recordings of sub-confluent LifeAct-mCherry expressing HUVECs (7-8 x 10^4^ cells/cm^2^) treated with 10 µM ROCK inhibitor Y27632 (overviews: scale bar 20 µm; cropped areas: 10 µm; time scale: mm:ss; compare Supplementary Movie 3). (I) Control (arrowheads indicate stress fibers). (II) After 3 minutes and (III) after 35 minutes following application of Y27632. Note the increased actin-driven protrusion after 3 minutes (crop 2, dotted lines), accompanied by loss of stress fibers and subsequent formation of junctional actin (crop 3, arrows). (b) Line scan analysis of LifeAct-mCherry intensities at t= 0 and t= 35 min after treatment with Y27632, as indicated in (a1, blue bar) and (a3, red bar) (**c**) SIM of confluent HUVECs immunolabeled as indicated under static conditions. Control cells (upper panel) and cells after treatment with Y27632 for 20 min (second panel). Cropped areas 1 and 2 (panels 3 and 4) are shown as indicated. Note the heterogeneous VE-cadherin and actin distribution with JAIL formation (crop 1, dotted lines) and the linearly organized VE-cadherin pattern associated with junctional actin formation (crop 2). Scale bar: 20 µm; crops 1 and 2: 5 µm. (**d**) Combined effect of U-FSS and Y27632 on HUVEC cultures. Cells were pre-incubated for 10 min with the ROCK inhibitor and then exposed to U-FSS for 20 min. Note the homogeneous linear VE-cadherin pattern with prominent overlap of actin and integrin β1 (arrowheads), compared to the corresponding DMSO control under shear stress. Scale bar: 20 µm. (**e, f**) Quantification of relative junctional (**e**) actin and (**f**) integrin β1 immunofluorescence intensity as indicated using the CBT (one dot= 15 cells; 3 independent experiments; one-way ANOVA). (**g**) Trans-endothelial electrical resistance (TER) determined in confluent HUVEC cultures as indicated. Note: U-FSS leads to characteristic TER changes, with an initial peak at t≈ 20 min followed by a subsequent decline, while Y27632 caused a rapid, transient decrease. Combined application of Y27632 and shear stress increased TER beyond the level observed with shear stress alone and thus counteracted the TER decrease induced by the ROCK inhibitor, resulting in an extended recovery phase and a sustained higher TER. (**h**) Quantitative data of the TER measurements. Bars shown represent mean values of maximum and minimum TER as indicated by “#”/“x” in (d). N = 4 independent experiments; unpaired Student’s t-test. (**i, k**) Phase-contrast time-lapse recordings of HUVEC cultures during TER measurements in the presence of Y27632 were further analyzed for apparent coherency using ImageJ. DMSO served as the control. (**k**) Statistical analysis of apparent coherency after t= 250 min compared to DMSO controls. Note the inhibitory effect on shear stress-induced adaptation, demonstrating that prolonged relaxation of actomyosin impairs the conversion process (unpaired Student’s t-test; n = 4 independent experiments). Error bars indicate ± SEM (ns β 0.05, *p < 0.05, **p < 0.01, ****p < 0.0001).

Functional assessment using trans-endothelial resistance (TER) measurements showed that Y27632 treatment rapidly decreased TER, reaching a minimum at ∼13 minutes and remaining below baseline for 2.5 hours (Figure 6d, for overview see Supplementary Fig.S5). This indicated compromised junctional integrity, consistent with irregular patterns of actin, integrin β1 and VE-cadherin.

To determine whether tension loss influences junctional integrity and barrier function in an agonistic or antagonistic manner during mechano-transduction, we next examined the effects of ROCK inhibition under shear stress conditions. Application of Y27632 under static conditions revealed a quick TER decreased, proofing the ROCK-inhibitor effect, which however, was totally counteracted by sudden onset of shear stress, reaching TER increased by about 30% compared to static control (Figure 6d). This obviously functional antagonistic effect is further confirmed by immunolabeling. Strikingly, a homogeneous linear VE-cadherin pattern combined with a prominent junctional actin and integrin β1 recruitment was observed together with a reduction of stress fibers (Figures 6e, 6g, 6h). These results demonstrate that shear stress-induced actomyosin tension loss complements, rather than opposes, the ROCK-mediated decrease in TER. This suggests that mechano-transduction physiologically reorganizes actin and VE-cadherin, likely through additional mechano-sensitive signaling pathways in addition to actomyosin relaxation. This aligns with the distinct dynamics and morphological patterns observed: homogeneous VE-cadherin linearization with recruitment of actin and integrin β1 after ROCK inhibition combined with shear stress (Figure 6e), whereas a more heterogeneous pattern occurred with ROCK inhibition under static conditions (Figure 6c). In addition, TER remained significantly higher in the presence of ROCK inhibition under shear stress than with shear stress alone for up to three hours, showing a clear delay in TER decrease. This suggests a delay in cell shape changes, as previously described ^15^. To investigate whether prolonged actomyosin relaxation affects shear stress-induced adaptation, experiments were performed in the presence of ROCK inhibition. Indeed, persistent relaxation reduced alignment and orientation after three hours of U-FSS (Figures 6i, 6k), indicating that a shift towards increased actomyosin tension after the intermediate state is necessary for physiological endothelial adaptation to shear stress. However, this transiently relaxed intermediate state may protect cells against overstimulation and structural damage, effectively priming them for adaptation.

### Adapted endothelial cells also transition through an intermediate state

The transient linearization and condensation of VE-cadherin, along with actin and integrin recruitment to cell junctions, defines a critical intermediate state required for physiological adaptation to shear stress. These findings raised the question whether the intermediate state is a common response to shifts in physiological shear stress or if it occurs exclusively in quiescent cells following onset shear stress. To investigate this, HUVEC cultures were first adapted to a shear stress of 6 dyn/cm^2^ for 6 hours, after which the shear stress level was increased to 36 dyn/cm^2^. Since we expected a complex morpho-dynamic response, we assessed both the aspect ratio using immunofluorescence snapshots (Figure 7a) and the continuous dynamic remodeling by determining the apparent coherency (Figure 7b). While AR is a simple, well-known parameter, apparent coherency is defined by two morphological parameters - cell shape and orientation - which can be simultaneously recorded using phase-contrast time-lapse recordings ^70, 71^. A value of 1 in apparent coherency reflects a uniform orientation in a dominant direction, while 0 indicates random orientation and round cell shape. Thus, we introduce apparent coherency as an novel metric for capturing the intricate relationship between shear stress-induced morphological changes and orientation in a single quantifiable value. Both parameters, AR (Figure 7a) and apparent coherency (Figure 7b), began to increase continuously within 1 hour after the onset of shear stress at 6 dyn/cm^2^ (Figures 7a (2), 7b (2)), and continued to rise over the next 12 hours (Figures 7a (5), 7b (5)). Notably, consistent with our hypothesis of an intermediate state during adaptation, a sudden increase in shear stress to 36 dyn/cm^2^ after 6 hours of adaptation caused a transient rise (Figures 7a (3), 7b (3)), followed by a sharp decrease in both AR (Figure 7a (4)), and apparent coherency within 2-3 hours (Figure 7b (4)). After this transient phase, both parameters steadily continued to increased (Figures 7a (6), 7b (6)).

**Figure 7.**
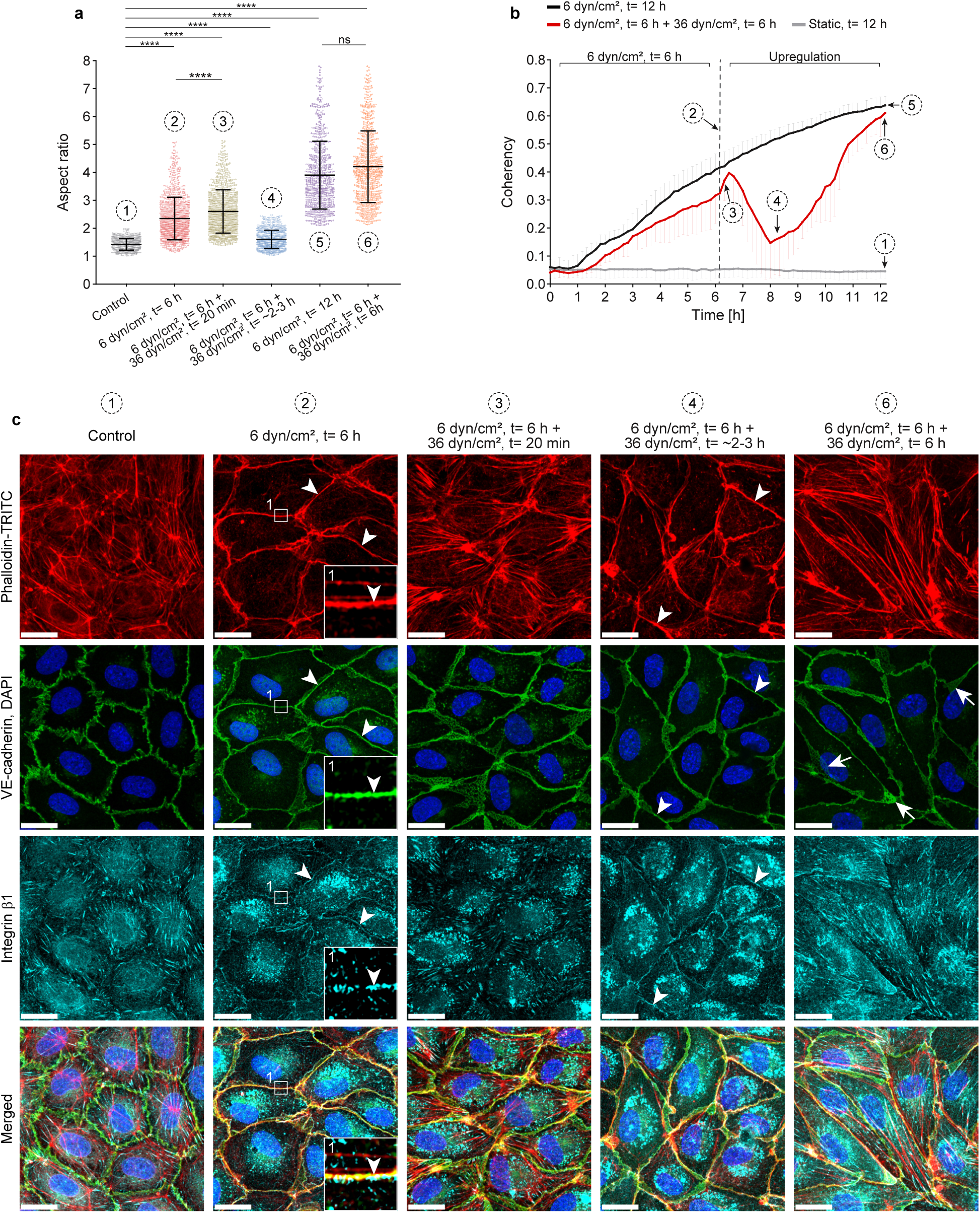
Adapted endothelial cells also transition through an intermediate state. Confluent HUVECs were exposed to different profiles of unidirectional fluid shear stress (U-FSS). Numbering (1-6) indicates different U-FSS profiles and is consistent throughout the sub-figures for comparison (**1**: static control; **2**: 6 dyn/cm^2^, t= 6 h; **3**: 6 dyn/cm^2^, t= 6 h + 36 dyn/cm^2^, t= 20 min; **4**: 6 dyn/cm^2^, t= 6 h + 36 dyn/cm^2^, t= ∼2-3 h; **5**: 6 dyn/cm^2^, t= 12 h; **6**: 6 dyn/cm^2^, t= 6 h + 36 dyn/cm^2^, t= 6 h). (**a**) Quantification of aspect ratio (cell elongation) from confluent HUVECs exposed to different U-FSS profiles as indicated using CBT based on immunofluorescence images (1: n = 784 cells; 2: n = 1107; 3: n = 1109; 4: n = 814; 5: n = 809; 6: n = 809; one-way ANOVA; error bars represent ± SEM; ns β 0.05, ****p < 0.0001). (**b**) Apparent coherency analysis of HUVECs exposed to U-FSS conditions as labelled using ImageJ OrientationJ plugin based on PhaCo time-lapse recordings. (Red line, number 4) Note the significant decrease in coherency approximately 2-3 h after upregulation to 36 dyn/cm^2^, followed by an accelerated increase (n ≈ 6500 cells/time point from 4 independent experiments). (**c**) Immunolabeling of HUVECs exposed to different U-FSS profiles analyzed by LSM as indicated. Scale bar: 20 µm; U-FSS direction from the left. (2) Note, after 6 dyn/cm^2^ for 6 h cells displayed a linear VE-cadherin pattern accompanied by a prominent junctional actin ring and integrin ß1 recruitment (arrowheads and SIM of cropped area 1). (3) Upregulation to 36 dyn/cm^2^ led to reappearance of stress fibers accompanied by cell elongation. (4) Note, after approximately 2-3 h of 36 dyn/cm^2^, cells regain a more cobblestone-like shape, characterized again by a linear VE-cadherin pattern, a prominent junctional actin ring and integrin β1 recruitment (arrowheads). (6) Upon prolonged exposure to 36 dyn/cm^2^, cells undergo reorientation and exhibit increased JAIL dynamics (arrows), actin stress fibers and elongation. In addition, compare Supplementary Figure 6 for enhanced overview images.

To elucidate the junctional organization of this complex remodeling, which is controlled by junctional dynamics ^21, 52^, fluorescent labelling of actin, VE-cadherin and β1-integrin was performed. Surprisingly, after 6 hours of flow at 6 dyn/cm^2^, the intermediate state was also observed and appeared prolonged, as most cells continued to exhibit a linear VE-cadherin pattern, junctional actin, and β1-integrin recruitment even after 6 hours (Figure 7c (2); overview in Supplementary Fig. 6(2)). However, a few cells displayed stress fibers and JAIL formation, indicated by VE-cadherin plaques. In addition, the prolonged intermediate state under 6 dyn/cm² was further confirmed, as the final adaptive response of the cells after 12 hours displayed the characteristic shear stress-induced architecture, with abundant stress fibers and large VE-cadherin plaques. This result is documented by overview images demonstrating the general and uniform cellular response under these conditions (Supplementary Fig. 6(5)). In contrast, the onset of 36 dyn/cm^2^ rapidly disrupted this prolonged intermediate state quickly within 20 minutes, as evidenced by loss of junctional actin, stress fibre re-formation, and increased VE-cadherin dynamics (Figure 7c (3); overview in Supplementary Fig. 6 (3)). However, intriguingly in already moderately adapted cells, the above-described intermediate state reappeared again after a sudden increase of shear stress to high levels of 36 dyn/cm^2^ within 2 hours. This renewed intermediate state is again characterized by prominent junctional actin with β1-integrin recruitment and VE-cadherin linearization (Figure 7c (4), overview in Supplementary Fig. 6 (4)). This effect strongly supports the broader relevance of an intermediate state along cell junctions for mechano-transduction-induced remodeling, which requires a cellular reset for this process. Following this intermediate state, adaptation to increased high shear stress accelerates, quickly leading to the loss of junctional actin, stress fiber formation, and VE-cadherin plaques typically observed previously (Figure 7c (6), Supplementary Fig. 6(6)). Consequently, these changes accelerate the adaptation, as shown by coherency analysis (Figure 7b (6)). Hence, the data confirm dose-dependent, shear stress-driven remodeling ^72^ with significant effects on junctional dynamics. Together, these findings demonstrate that the intermediate state - defined by transient VE-cadherin linearization, junctional actin, and β1-integrin recruitment - is a fundamental and recurring feature of endothelial adaptation to changes in physiological shear stress, rather than a response limited to quiescent cells.

## Discussion

The transition of endothelial cells from a resting (quiescent) to an active state and vice versa is a fundamental process in vascular biology, underlying responses to hemodynamic forces, inflammation, and tissue remodeling ^7, 14, 17, 53^. Resting endothelial cells represent the final stage during tissue formation and in cell culture. They maintain the vascular barrier and regulate homeostasis by controlling intercellular junctions and cell-substrate contacts ^73–76^. In contrast, activating stimuli such as shear stress, wound healing, or inflammatory signals prompt a shift toward a dynamic, remodeling phenotype ^7–10, 18, 35, 77^. This shift involves rapid cytoskeletal reorganization, changes in junctional protein distribution, and upregulation of adhesion molecules, enabling increased cell migration, proliferation, and adaptation to new functional demands. Understanding the molecular and cellular mechanisms governing this transition is crucial for elucidating how endothelial cells balance stability with plasticity in both physiological adaptation and disease contexts.

### Mechanistic interplay between cell circumference, junctional VE-cadherin concentration and actin dynamics during endothelial maturation

Our findings demonstrate that the transition from an active, polymorphic endothelial phenotype to a quiescent, cobblestone-like monolayer during growth is accompanied by significant changes in cell size and dynamics, including a reduction in cell circumference. The establishment of this stable monolayer in response to increasing cell density involves coordinated remodeling of VE-cadherin, actin, and integrins. We show that linearly organized VE-cadherin clustering coincides with the emergence of prominent linear junctional actin filaments and integrins, together with the loss of actin stress fibers, all hallmarks of junction maturation. Notably, the reduction in cell circumference increases the local concentration of junctional VE-cadherin, which in turn suppresses VE-cadherin-mediated dynamics, particularly JAIL, thereby promoting junctional stability. This is consistent with our observation that total VE-cadherin and actin expression remain unchanged in HUVEC culture models and during angiogenesis *in vivo* ^21, 48, 52, 78^. Furthermore, the observed decrease in G-actin levels with increasing cell density, while total actin levels remain stable, supports the notion that junction maturation primarily involves a reorganization of both VE-cadherin and actin rather than new synthesis. In addition, the mature phenotype is in line with a resting endothelium in large vessels ^54, 55, 79^. Strong mechanistically evidence for the critical role of junctional VE-cadherin concentration for maturation is further demonstrated by overexpressing VE-cadherin in sub-confluent cell layers (7 x 10^4^ cells/cm^2^). Although these cells exhibit stress fibers and irregular actin, integrin, and VE-cadherin distribution, VE-cadherin overexpression induced all the characteristics of a mature monolayer, including prominent junctional actin colocalized with integrins and VE-cadherin. Functionally, this resulted in reduced cell junction and overall cell dynamics. Collectively, these results highlight the interdependence of cell size and the critical role of junctional VE-cadherin concentration in determining actin-driven cell dynamics during the maturation and stabilization of endothelial junctions.

### The shift from quiescent to active endothelium involves a protective, tension-sensitive intermediate state

Endothelial cells are well known to change their phenotype in response to shear stress through various mechano-transduction mechanisms ^14, 15, 60, 80^. Here, we applied fluid shear stress as a stimulus to convert a largely mature endothelial monolayer toward an arterial phenotype and to study the associated changes in actin-filaments, cell junctions and cell-substrate interactions. We demonstrate that this conversion consistently involves an intermediate state characterized by temporary stabilization of cell junctions. This effect, occurring rapidly within 20 minutes after the onset of shear stress, is described here for the first time and provides new insights into this complex remodeling process. The intermediate state is marked by a transient increase in trans-endothelial resistance, with its morphological correlate in VE-cadherin clustering and actin recruitment, which in turn targets integrins to these sites and stabilizes cell adhesion complexes, as visible already by phase-contrast microscopy. Remarkably, these changes in the molecular organization of the cellular phenotype resemble a mature cell monolayer and is observed after both the onset and upregulation of shear stress in already shear stress adapted cell monolayers as well as after VE-cadherin overexpression.

Mechanistically, onset of the intermediate state was due to the dephosphorylation of myosin light chain (MLC), a critical regulator of barrier function and contractility ^67, 81^. This shear stress-induced, dephosphorylation drives junctional actin recruitment, as further supported by the combined application with a ROCK inhibitor-dependent relaxation. This effect of ROCK inhibition has also been shown to preserve tension-dependent junctional integrity ^82^. In our approach, the dephosphorylation-dependent tension loss was confirmed by laser ablation experiments, Western blotting, and immunolabeling, and is consistent with shear stress-induced tension loss demonstrated using a tension sensor coupled to VE-cadherin ^80^.

Considering its distinct functional properties, it is plausible to envision that the stress-induced reversion to a mature-like phenotype acts as a mechanistic reset, serving as a protective measure that shields the endothelium from sudden shear stress-induced damage while simultaneously allowing controlled and precise remodeling. Further support for this idea comes from observations that this intermediate state emerges following abrupt increases in shear stress in already adapted cells, and that the shear stress-induced, transient increase in barrier function shows dose dependency ^72^. Similar effects have also been reported after challenging HUVECs with VEGF, which, as observed here upon shear stress, are accompanied by a transient increase in barrier function and transient MLC dephosphorylation ^9, 83^. Moreover, VE-cadherin linearization and actin recruitment have been observed in TNF-α-stimulated cells, accompanied by a transient increase in barrier function ^35^. Taken together, these observations suggest that the intermediate state is part of a broader, integrated cellular regulatory framework that enables a general response to stress. This process may also prime genetic and epigenetic mechanisms that support further adaptive responses in both physiological and pathological contexts.

### VE-cadherin as a central regulator of endothelial state transitions

Our data indicates that VE-cadherin acts as a key determinant in endothelial remodeling processes. This includes the maturation phase, during which VE-cadherin increasingly accumulates at cell junctions, thereby stabilizing them, reducing junctional actin dynamics, and promoting a resting phenotype with high barrier function. In contrast, the transition from a resting to an active phenotype requires VE-cadherin dilution. This parameter is simple controlled by cell shape changes that increase the cell circumference, thereby diluting the available junctional VE-cadherin. In line with this, VE-cadherin overexpression largely blocked shear stress-induced long-term adaptation, as well as previously described processes such as wound healing and angiogenesis, by suppressing overall cell junction dynamics (for review see ^21^). The critical role of VE-cadherin in remodeling is further underscored by its concerted interaction with actin and integrins, indicating a central regulatory function at both intercellular junctions and cell-substrate interfaces. This is supported by observations that VE-cadherin overexpression in sub-confluent cells directs actin filament recruitment and, in turn, integrin localization, resulting in a phenotype resembling that of high-density cultures with physiologically mature junctions. Together, these findings strongly support the concept that VE-cadherin is a key determinant in both endothelial maturation and the transition to an active state. Furthermore, we show that actin is recruited to cell junctions when VE-cadherin accumulates - both during the maturation phase in highly confluent cells and following VE-cadherin overexpression in sub-confluent cells - and that this is accompanied by the recruitment of integrin β1 and integrin αvβ3 to the same sites. This process depends on tension modulation regulated by MLC phosphorylation. The interplay between integrins and VE-cadherin has been shown to be important for both angiogenesis ^20^ and barrier function ^84^. These studies highlight a critical role for integrins in organizing junctional VE-cadherin, as integrin depletion disrupts VE-cadherin organization, which appears contradictory to the concept that VE-cadherin recruits actin and, in turn, integrins. However, considering the interplay of actin with both integrins and VE-cadherin, this apparent contradiction is challenged. Notably, there are reports demonstrating actin-mediated stabilization of both VE-cadherin complexes and integrin complexes, as well as evidence for the reverse relationship ^22, 41, 85–87^.Therefore, it should be considered that the dynamic behavior of both cell-cell junctions and cell-substrate adhesions is determined by the coordinated interplay of VE-cadherin, actin, and integrins, rather than by any single cytoskeletal or adhesion molecule alone. Based on our data, we propose that junctional actin provides the physical link between the VE-cadherin/catenin complex and integrin-mediated cell-substrate adhesions. The combined binding of junctional actin by the VE-cadherin/catenin complex and cell-substrate adhesion complexes provides robust stabilization of the cell monolayer. The junctional regulation of the VE-cadherin/catenin complex, actin, and integrins has been shown to involve Rho GTPases, tyrosine phosphorylation signals, and additional components of cell-cell and cell-substrate adhesion complexes, such as VE-PTP and focal adhesion kinases ^43, 88, 89^. Understanding how these additional control mechanisms coordinate the combined interplay between cell junctions and cell-substrate adhesions warrants further investigation.

In summary, our findings show that junctional VE-cadherin levels, actomyosin-mediated tension control, and actin dynamics, together with integrins, coordinate to regulate endothelial cell state transitions. The conversion from a resting to an active state includes a tension-sensitive protective intermediate state, which extends the current mechanistic understanding of how endothelial cells maintain junction stability and undergo remodeling. This mechanism critically depends on JAIL-driven junction dynamics, which provide the mechanical force for remodeling and enable precise control of barrier function by modulating its dynamics during this process. These insights highlight how adhesion, force balance, and structural adaptation are tightly coordinated during cellular maturation and the transition to an active phenotype. An intermediate cellular state that emerges during this transition appears to confer protection against stress and may reset cells for controlled remodeling. This provides a general framework for understanding vascular function, tissue maintenance, and cellular responses to stress, as illustrated in Figure 8 (Figure 8).

**Figure 8.**
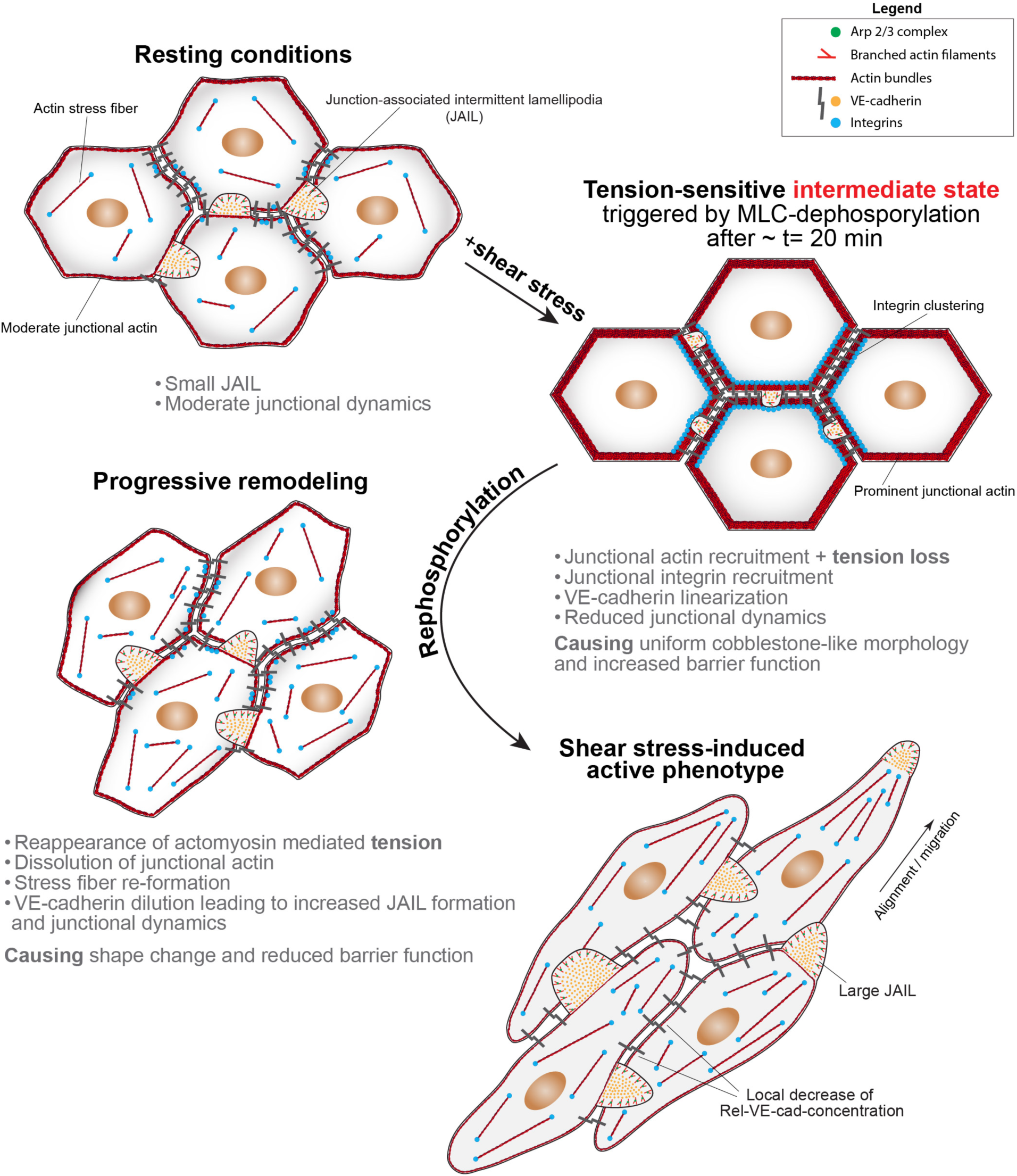
Scheme illustrating the conversion of resting cells to an active state using physiological shear stress. Resting HUVEC cultures used for conversion to an active phenotype had a cell density of 10 x 10^4^ cells/cm^2^, which led to moderate junctional actin organization and high VE-cadherin concentration, with low junctional dynamics indicated by small junction-associated intermittent lamellipodia (JAILs). The onset of shear stress induced an intermediate state triggered by transient myosin light chain (MLC) dephosphorylation. This mechanism caused prominent recruitment of junctional actin together with integrins (shown for β1 and α_V_β3), linearization of VE-cadherin, increased barrier function, and further downregulation of junction dynamics, as indicated by very small JAILs, resulting in a uniform cobblestone-like morphology. This intermediate state was a general phenomenon also observed in cells already adapted to shear stress. Subsequent MLC re-phosphorylation initiated the transition to an arterial phenotype by dissolving junctional actin, leading to stress fiber reformation and VE-cadherin dilution, which increased junctional dynamics through JAIL formation, driving changes in cell shape and promoting migration. This process was accompanied by a decrease in barrier function due to cell junction weakening, which is required for remodeling. Ultimately, the cells developed an elongated, shear stress-aligned phenotype, as frequently described.

## Materials and Methods

### Cell culture

Human umbilical vein endothelial cells (HUVECs) were isolated from umbilical cord veins of different donors (according to ^9^), a procedure approved by the ethics committee of the University of Muenster (2009-537-f-S): briefly, umbilical cord veins were perfused with PBS (Thermo Fisher Scientific, Waltham, USA) and then digested with 1 mg/ml collagenase (Biochrom, Berlin, Germany) diluted in PBS for 10 min at 37 °C in a water bath. Subsequently, detached cells were collected and centrifuged at 1200 rpm for 5 min. ECs were grown in endothelial growth medium (PromoCell, Heidelberg, Germany) supplemented with endothelial cell growth medium mix and 1% penicillin/streptomycin (P/S) on gelatin-coated (Sigma-Aldrich, St. Louis, USA) tissue culture flasks at 5% CO_2_ and 37 °C. For all experiments HUVECs from passage 0 or 1 were used. HEK 293T and HEK 293 cells, provided by the American Type Culture Collection (Manassas, USA), were maintained in high-glucose DMEM (Sigma-Aldrich, St. Louis, USA) supplemented with 10% fetal calf serum (FCS), 1% P/S, and 1% glutamine.

### Crosslinked gelatin coating procedure

In all experiments, cells were seeded on crosslinked gelatin-coated and bleached surfaces. Surfaces were coated with 0.5% gelatin (Sigma-Aldrich, St. Louis, USA) for 30 min at 37 °C and then crosslinked with 2% glutaraldehyde (Sigma-Aldrich, St. Louis, USA) for 8 min. After washing, 70% ethanol was added for 1 h at room temperature (RT), followed by four washing steps with PBS. Subsequently, UV bleaching was performed for 45 min (Biometra, Göttingen, Germany). Finally, chambers were washed with PBS and incubated in 2mM glycine/PBS + 1% ascorbic acid overnight at RT. Before cell seeding, chambers were washed four times with PBS.

### Immunofluorescence staining of cell cultures

After experiments, endothelial cells were gently rinsed with prewarmed M199 medium (Sigma-Aldrich, St. Louis, USA), followed by fixation with 4% paraformaldehyde/PBS, pH 7.3 for 10 min and washed three times with PBS. HUVECs were then permeabilized in PBS/0.5% Triton X-100 (Thermo Fisher Scientific, Waltham, USA) for 10 min at RT, rinsed three times for 5 min each in PBS/1% BSA, subsequently blocked in antibody diluting solution (PBS, 0.1% Triton X-100, 2% BSA) at RT for 1h and incubated overnight in antibody diluting solution at 4 °C with goat anti-VE-cadherin (1:200, #AF938, R&D Systems, Minneapolis, USA), rabbit anti-ß1 integrin (1:1000) ^90^, mouse anti-a_V_ß3 integrin (1:400, #MAB1976, Millipore, Burlington, USA), rabbit anti-phospho myosin light chain II (ser19) (1:100, #3671, Cell Signaling Technology, Danvers, USA) and mouse anti-beta-catenin (1:200, #610154, BD Biosciences, USA). After four 5-min washes in PBS/1% BSA, samples were incubated in PBS/1% BSA for 1 h at RT with corresponding Alexa Fluor conjugated secondary antibodies (1:200, Invitrogen, Carlsbad, USA: anti-goat 633; anti-rabbit 488; anti-mouse 488; anti-mouse 647), Phalloidin-TRITC (1:400, Sigma-Aldrich, St. Louis, USA) and Palloidin-647 (1:100, #ab176759, Abcam, Cambridge, UK). Finally, cells were rinsed three times with PBS/1% BSA, stained with DAPI (1:10000, #28718-90-3, Sigma-Aldrich, St. Louis, USA) and mounted in DAKO fluorescent mounting medium (#S3023, Agilent Technologies, Santa Clara, USA).

### Microscopy

An Axio observer Z1 microscope (Carl Zeiss, Oberkochen, Germany) equipped with a 37 °C stage, 37 °C flow unit (BioTechFlow system (BTF system), MOS Technologies, Telgte, Germany) and 5% CO_2_ supply was used with Zeiss A-Plan 10x/0,25 Ph1 M27 objective for phase contrast live cell imaging. Laser scanning microscopy (LSM), Structured illumination microscopy (SIM) and Total internal reflection fluorescence (TIRF) microscopy was conducted by using a LSM 780 confocal microscope complemented with the ELYRA super resolution module with 405, 488, 561, and 647 nm diode lasers (Carl Zeiss, Oberkochen, Germany). LSM and SIM images were acquired using Zeiss objectives: EC Plan-Neofluar 40x/1.30 Oil M27 or Plan-Apochromat 63x/1.4 Oil DIC M27 (Carl Zeiss, Oberkochen, Germany). For TIRF imaging, a Zeiss alpha Plan-Apochromat 100x/1.46 Oil DIC M27 objective was used. SIM images were processed for reconstruction with ZEN imaging software (Carl Zeiss, Oberkochen, Germany). A spinning disk confocal microscope (SpDM) (Carl Zeiss, Oberkochen, Germany) equipped with a single BTF unit (MOS Technologies), definite focus, 37 °C heated stage, humidity and 5% CO_2_ supply was used for VE-cadherin-EGFP and LifeAct-mCherry fluorescence live cell imaging. High temporal resolution fluorescence time-lapse imaging was performed every 10-20 sec on HUVECs subjected to shear stress or treated with ROCK inhibitor Y27632 (10 µM, Sigma-Aldrich, St. Louis, USA) using a Zeiss Plan Apochromat 40x/1.4 oil DIC (UV) VIS-IR immersion objective (Carl Zeiss, Oberkochen, Germany). The same acquisition settings were consistently applied to all samples within the same biological replicate. Image processing, brightness and contrast were adjusted with Fiji ImageJ (National Institutes of Health, USA).

### Quantification of total actin in sub-confluent and confluent HUVEC cultures

Seeding the same number of cells into cell culture dishes of different diameters (35 mm and 85 mm). When cells in the smaller dish reached confluence ([9-10] x10^4^ cells/cm^2^), the cell culture in the larger dish was still sub-confluent ([7-8]x10^4^ cells/cm^2^). Cells were lysed and equal amounts of total cellular protein from sub-confluent and confluent cell lysates (5 µg, 10 µg and 20 μg) were determined by amido-black method ^91^, transferred to SDS gel electrophoresis, prepared for quantitative western blotting and probed with mouse anti-pan-actin antibody (1:1000, #sc-56459, Santa Cruz Biotechnology, Dallas, USA) and mouse anti-alpha tubulin (1:5000, #B-5-1-2, Sigma-Aldrich, St. Louis, USA). The mean intensity of the normalized values was plotted as a function of total protein amount. Subsequently, linear regression analysis was done utilizing GraphPad PRISM software (GraphPad Software, version 10, La Jolla, USA).

### One-drop assay

The drop assay was established to study cell populations of different densities (sub-confluent (7-8 x 10^4^ cells/cm^2^), confluent (9-10 x 10^4^ cells/cm^2^) and highly confluent (11-12 x 10^4^ cells/cm^2^)) within the same cell culture and was previously described ^22^. Briefly, a 20-30 µl drop of approximately 8 x 10^3^ - 1 x 10^4^ cells was initially added to the center of a crosslinked gelatin coated glass bottom (#1.5) cell culture dish and incubated for 1 h at 5% CO_2_ and 37 °C. Afterwards, cell medium was added and HUVECs were cultured for 6-8 days. This leads to cell accumulation in the central area, while cell density decreases with increasing distance from the center (compare scheme in Supplementary Fig. 1b).

### G-actin assay

HUVECs were seeded onto a crosslinked gelatin-coated glass bottom dish using the one-drop assay technique, enabling the analysis of both sub-confluent and confluent cell populations within the same culture. G-actin immunostainings were performed according to the protocol described ^92^: briefly, cells were fixed for 20 minutes in 4% paraformaldehyde (PFA) solution supplemented with 10 mM MES, 3 mM MgCl_2_, 138 mM KCl, 2 mM EGTA, 0.32 M sucrose and adjusted to pH 6.1. After washing with PBS, HUVECs were permeabilized in PBS/0.5% Triton X-100 (Thermo Fisher Scientific, Waltham, USA) for 10 min, rinsed in PBS/0.1% Triton X-100, subsequently blocked in antibody diluting solution (PBS, 0.1% Triton X-100, 2% BSA) at RT for 1 h and incubated overnight at 4 °C with goat anti-VE-cadherin antibody (1:200, #AF938, R&D Systems, Minneapolis, USA) for cell border detection. After washing with PBS/1% BSA, samples were incubated for 30-45 min at RT in antibody diluting solution with anti-goat Alexa Fluor 633 antibody (1:200, Invitrogen, Carlsbad, USA), Phalloidin-TRITC (1:400, Sigma-Aldrich, St. Louis, USA) and 2 µg/ml Alexa 488 conjugated deoxyribonuclease I (DNase I, #D12371, Thermo Fisher Scientific, Waltham, USA) for G-actin labeling. Finally, cells were rinsed three times with PBS/1% BSA, mounted in DAKO fluorescent mounting medium and Z-stack projections were acquired by using a LSM 780 microscope (Carl Zeiss, Oberkochen, Germany). Cells were segmented based on the VE-cadherin stainings with the CellBorderTracker (CBT) ^93^, which allowed the measurement of G-actin fluorescence intensity per cell and cell perimeter by using Fiji ImageJ software (National Institutes of Health, USA).

### Quantification of cell morpho-dynamics and junctional fluorescent intensities

Cell migration quantification was conducted by manual cell tracking utilizing the Fiji ’Manual Tracking’ plugin (https://imagej.nih.gov/ij/plugins/track/track.html) alongside the ’Chemotaxis and Migration Tool’ based on phase contrast time-lapse recordings. The analysis involved the comparison of several parameters, in particular: i) cell velocity, ii) accumulated distance, representing the total path length traveled by the cell, and iii) Euclidean distance, indicating the direct distance between the initial and final points of cell movement.

Apparent coherency analysis ^70, 71^ of HUVECs exposed to different shear stress profiles was performed using the ImageJ ’OrientationJ’ plugin and the ’Dominant Direction’ tool based on phase contrast time-lapse images. Phase contrast images were acquired at 10x magnification (1421 µm x 1065 µm) and adjusted for coherency analysis. Apparent coherency, ranging from 0 to 1, quantifies the uniformity of cell shape and orientation. A value of 1 indicates strong, uniform alignment in a dominant direction, whereas a value of 0 reflects random orientation. This ideal metric combines cell elongation and overall shape to quantify consistent cell orientation.

The CellBorderTracker (CBT) software ^93^ was employed to determine the aspect ratio (ratio between the major and minor axis of an ellipse fitted to the outline of each cell) and perimeter of fixed cells. For this, cells were segmented based on VE-cadherin staining and transferred to Fiji ImageJ (National Institutes of Health, USA), where morphological parameters were quantified using the ’Analyze Particle’ function. The fluorescence intensities of actin, integrin ß1 and integrin a_V_ß3 at the junctions were also quantified using the CBT. Maximum intensity projections of fluorescence images at 63x magnification (135µmx135µm) were uniformly used for these quantifications. After segmentation, Fiji ImageJ (National Institutes of Health, USA) was utilized to generate a distance map for defining the region of interest (ROI) at the cell junctions. The ROI was defined by extending 5 pixels on each side of the segmentation line. The relative concentration of junctional actin, integrin ß1 and integrin a_V_ß3 was defined as the integrated intensity of the respective proteins in the ROI divided by the total cell length of the ROI (compare Supplementary Fig. 2a). Furthermore, for shear stress experiments, the aspect ratio was automatically calculated using a custom-designed MATLAB (MathWorks, Natick, MA, USA) based on high-resolution phase-contrast live-cell imaging.

### Measurement of VE-cadherin-EGFP displacement

To estimate junctional dynamics, time-lapse movies of VE-cadherin-EGFP expressing cells were segmented using the CellBorderTracker ^93^. From the resulting segmentation, the overall distance (d-mean) of the cell-borders at time point t and time point t-dt as well as time point t+Δt was estimated using a self-written MATLAB script. The average junctional dynamic displacement at time point t can then be estimated by d-mean(t)/(2xΔt).

### Plasmid construction and lentivirus production

The detailed procedure for the generation of fluorescence-tagged VE-cadherin-EGFP and LifeAct-mCherry and their cloning into the lentiviral pFUGW vector was recently outlined in ^36, 52^. Lentivirus was generated in 293T cells grown in DMEM (high glucose plus 10% FCS and 1% P/S) medium in 15 cm cell culture dishes. Cells were transfected with the pFUGW vector containing the gene of interest, along with the packaging vector pCMV-ΔR8.74 and the envelope vector pMD2G, which carries the VSV glycoprotein. Specifically, the plasmids pFUGW with the gene of interest (23 μg/dish), pCMV-ΔR8.74 (23 μg/dish), and pMD2G (11.5 μg/dish) were dissolved in 1725μl of DMEM medium without FCS and antibiotics (DMEM-/-). The transfection mixture, which included 172.5μl of polyethylenimine transfection reagent (1 mg/ml, Polysciences, USA), was dissolved in 1600.8 μl of DMEM-/- and incubated at RT for 30 minutes. This mixture was then combined with the plasmid solution and transfected into 293T cells. Following a 15-18 hours incubation period at 37°C, the medium was replaced with fresh, complete DMEM medium supplemented with 10% FCS, 1% penicillin/streptomycin (P/S) and 1% glutamine. After 24 hours, the supernatant was collected, centrifuged at 3000 rpm for 10 minutes, and sterile-filtered using a 0.45μm filter. Lentivirus particles were then concentrated via ultracentrifugation at 25000 rpm for 1.5 hours at 4°C (Beckman Coulter ultracentrifuge, Palo Alto, USA). The pellet was resuspended in 150μl of PBS containing 1% BSA and incubated for 1 hour at 4°C. Aliquots were subsequently stored at -80°C. Before use in experiments, the virus was titrated and tested in a bioassay using HUVEC cell cultures. For subsequent transduction experiments, the lowest virus concentration that provided a sufficient signal was selected.

### Adenovirus production

For VE-cadherin-EGFP overexpression, adenovirus containing VE-cadherin-EGFP or EGFP (kindly provided by Masatoshi Takeichi, ^94^ was used and amplified as described below. HEK 293 cells were seeded in a tissue culture flask and grown to 80-90% confluence. The medium was changed to DMEM (Biochrom, Berlin, Germany) without FCS and P/S supplemented with 1% glutamine and 10μL (15 MOL) adenovirus stock. After incubation for 1 hour at 37°C, equal volumes of DMEM containing 4% FCS and 1% glutamine were added to achieve a final concentration of 2% FCS in the cell culture medium. After 2 days, four cycles of freeze-thaw-vortex were performed to harvest the virus. Cells were removed from the plate and centrifuged at 500 x g for 10 min. The pellet was then resuspended in 6 ml PBS, followed by four rounds of freeze-thawing and centrifugation at 4000 x g for 20 min at 4°C.

The supernatant was vortexed with 3.3 g of ultrapure CsCl and centrifuged at 1.76 × 10^5^ x g for 22 h at 10°C. Subsequently, virus was harvested using glycerol (10%), divided into aliquots, and stored at -80°C. The effect of overexpression has been previously documented elsewhere by quantitative Western blot ^9, 35^.

### Gene transduction

For lentiviral-mediated gene transduction, 2 x 10^4^ cells/cm^2^ were initially seeded in the afternoon. After 15-18 hours, the medium was aspirated from the cells, collected and replaced with the viral particle mix dissolved in PromoCell medium (without FCS and P/S) (PromoCell, Heidelberg, Germany) supplemented with 3% polyvinylpyrrolidone (PVP) (Sigma-Aldrich, St. Louis, USA) for 1 hour at 37°C. PVP enhances transduction efficiency. The previously collected supernatant was then returned to the cells, and fresh PromoCell medium containing FCS and P/S was added the following day. The medium was subsequently changed every two days until the experiment was conducted. For adenovirus-mediated gene transduction, HUVECs were transduced with an adenoviral mix in PromoCell medium (without FCS and P/S) supplemented with 3% PVP on the evening before the experiment. After a 4-hour incubation at 37°C, the virus mixture was aspirated and replaced with fresh PromoCell medium.

### Laser Ablation

Nanoablation was performed in LifeAct-mCherry expressing HUVEC cultures according to a method previously described^95^. Briefly, an Observer Z1 microscope (Carl Zeiss, Oberkochen, Germany) with a Yokogawa CSU10B spinning disk and sCMOS ORCA Flash 4.0LT system (Yokogawa Electric, Tokyo, Japan) plus a Zeiss alpha Plan-Apochromat 100x/1.46 Oil DIC (UV) VIS-IR objective (Carl Zeiss, Oberkochen, Germany) was used. The microscope was additionally equipped with a DPSL-355/14 ultraviolet laser ablation system (Rapp OptoElectronics, Wedel, Germany). Following laser calibration, the laser power was reduced to 2% for all experiments. Time-lapse images were captured every 200 ms to track LifeAct-mCherry displacement behavior after the laser cut. Recoil velocity and displacement were analyzed using the line scan and kymograph tools in Fiji ImageJ. For further quantification, the well-established Kelvin-Voigt mathematical-physical model was applied to determine the elastic stiffness tau (τ) of junctional actin, serving as an indicator of tension ^96–98^. The time-lapse data were fitted to the following exponential function using MATLAB’s Curve Fitting tool (MathWorks, Natick, MA, USA) to determine tau (τ):

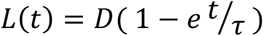

L(t) describes the displacement over time t and D indicates the force/elasticity ratio. Higher tension is indicated by decreased τ levels and less tension by increased τ.

### Western blotting

Cells were lysed in SDS sample buffer (0.18 M Tris, 6% SDS, 30% glycerine, 80 mg/l Bromophenolblue, 10 mM DTT). The total protein concentration of the samples was determined by the Amido Black method prior to gel loading ^91^. Thus, each lane of a polyacrylamide gel was loaded with equal amounts of proteins. After gel electrophoresis, western blotting was performed and wet transferred to a 0.45 µm PVDF membrane. Membranes were incubated in PBST/5% skim milk with mouse anti-α-tubulin (1:5000, #T5168, Sigma-Aldrich, St. Louis, USA), mouse anti-EGFP (1:1000, #632569, Takara Bio Clontech, Shiga, Japan), goat anti-VE-cadherin (1:800, #AF938, R&D Systems, Minneapolis, USA), rabbit anti-phospho myosin light chain II (ser19) (1:500, #3671, Cell Signaling Technology, Danvers, USA), mouse anti-pan actin (1:1000, #sc-56459, Santa Cruz Biotechnology, Dallas, USA), mouse anti-ß1 integrin (1:1000, #ab30394, Abcam, Cambridge, UK) overnight at 4°C. Afterwards, protein bands were labeled for 1 h at RT using IRDye 680 RD and 800 RD infrared secondary anti-mouse, anti-goat and anti-rabbit antibodies (1:10000, LICOR Biosciences, Bad Homburg, Germany) and detected by Li-Cor Odyssey Infrared Reading System (LICOR Biosciences, Bad Homburg, Germany). The Odyssey software package (Li-Cor) was employed for quantitative analysis.

### Shear stress experiments

Defined levels of unidirectional fluid shear stress (indicated in figures) were generated in the BioTechFlow system, a cone and plate in vitro rheological system (MOS Technologies, Telgte, Germany) as described in ^58, 59^. Briefly, HUVECs were initially seeded at a density of 3 x 10^4^ cells/cm^2^ on BTF-glass plates (27 cm^2^) coated with crosslinked gelatin and mounted in stainless steel flow chambers. Cells were cultured to confluence over 4-5 days. During flow experiments, endothelial cells (ECs) were maintained in PromoCell medium supplemented with 3% PVP (Sigma-Aldrich, St. Louis, USA) to increase viscosity, at 5% CO₂ and 37 °C. A transparent rotating cone (2.5°) was positioned above the cell layer, with its tip nearly touching the endothelial cell surface. The cone’s rotation generated a stable three-dimensional laminar flow, enabling continuous medium exchange above the EC cultures. Since the entire cone-plate system is transparent, both fluorescent microscopy analysis of fixed samples post-flow and phase-contrast time-lapse imaging during shear stress exposure are feasible. To achieve high temporal resolution, phase-contrast and fluorescent live cell time-lapse images were captured every 10-20 seconds. Fluorescence live-cell imaging under shear stress was conducted using SpDM as previously described. HUVECs expressing VE-cadherin-EGFP or LifeAct-mCherry were cultured on a round coverslip (65 mm, #1.5), which was placed in a custom-made BTF chamber to prevent deformation of the delicate slides during shear stress application.

### Impedance measurements under shear stress conditions

Trans-endothelial electrical resistance (TER) of confluent HUVEC monolayers under unidirectional fluid shear stress was determined by using custom-designed impedance chambers^15^. Cells were prepared as described above and exposed to a maximum of 18 dyn/cm^2^ U-FSS. For ROCK inhibition, ROCK inhibitor Y27632 (10 µM, Sigma-Aldrich, St. Louis, USA) was used. Prior to flow, cells were incubated with Y27632 for 10 minutes or with the appropriate concentration of dimethyl sulfoxide (DMSO) (Sigma-Aldrich, St. Louis, USA) in the controls. Impedance was determined every twelve seconds using an impedance generator and analyzer (National Instruments, USA). Data were subsequently analyzed with a custom-developed TER analytical software (LabView, National Instruments, USA) from MOS Technologies (Telgte, Germany) and calculated as TER(t)/TER(t=0), representing the TER at a given time (t) divided by the TER at time zero. Time 0 is defined as the time point of treatment with Y27632 or DMSO. Experiments and controls were conducted simultaneously using the fully automated BTF system with 9 individual units (MOS Technologies, Telgte, Germany).

### Statistical analysis

The statistical analyses were performed using GraphPad PRISM software (GraphPad Software, version 10, La Jolla, USA). To assess normality, the data were analyzed using the D’Agostino-Pearson and Shapiro-Wilk test. An unpaired two-tailed Student’s t-test was used to compare two groups with normally distributed data and Mann-Whitney test was used for non-normally distributed data. For comparisons between more than two groups, a one-way ANOVA test was applied. Spearman correlation coefficient was used to evaluate correlation between G-actin intensity and cell perimeter. Differences were considered significant when p < 0.05 (*p < 0.05; **p < 0.01; ***p < 0.001; ****p < 0.0001). Unless otherwise specified, all data were obtained from at least three independent experiments.

## Acknowledgements

We thank Annelie Ahle and Christine Schimp for their skillful technical assistance. Masatoshi Takeichi generously provided us with VE-cadherin-EGFP adenoviral constructs. We are grateful to Till Rauterberg for his editing and help in preparing the movies. Special thanks to Dietmar Vestweber for critical discussions on the topic. This work was supported by grants from the Deutsche Forschungsgemeinschaft (DFG) and the BMBF (03ZZ0902D) awarded to H. S. (SCHN 430/9-1 and INST 2105/24-1), from the DFG awarded to E. R. (CRC1348 B06) and M. G. (GA 2268/4-1), from the Max Planck Society awarded to M. O.-S. and H. S., and from the *Medizinerkolleg* (MedK) of the Medical Faculty Münster awarded to J. F. We are grateful for the generous support of Werner Paulus and colleagues at the Institute of Neuropathology, University Hospital of Münster.

## Author contributions

Author contributions J.F. performed the majority of the experiments, analyzed the data and prepared the figures with significant help of M.O-S. J.P.K. and V.B. contributed to overexpression experiments. F.M. performed migration assays and J.S assisted in the analyses. Z.L and E.R helped with laser ablation experiments. M.G. provided expert advice on coherency analyses. J.E. provided expert advice to integrins. H.S. initiated the topic, supervised the entire work, and wrote the manuscript together with J.F. All authors contributed with discussions, suggestions and critical reading and editing the MS.

## Supplementary figures

**Supplementary Figure 1:**
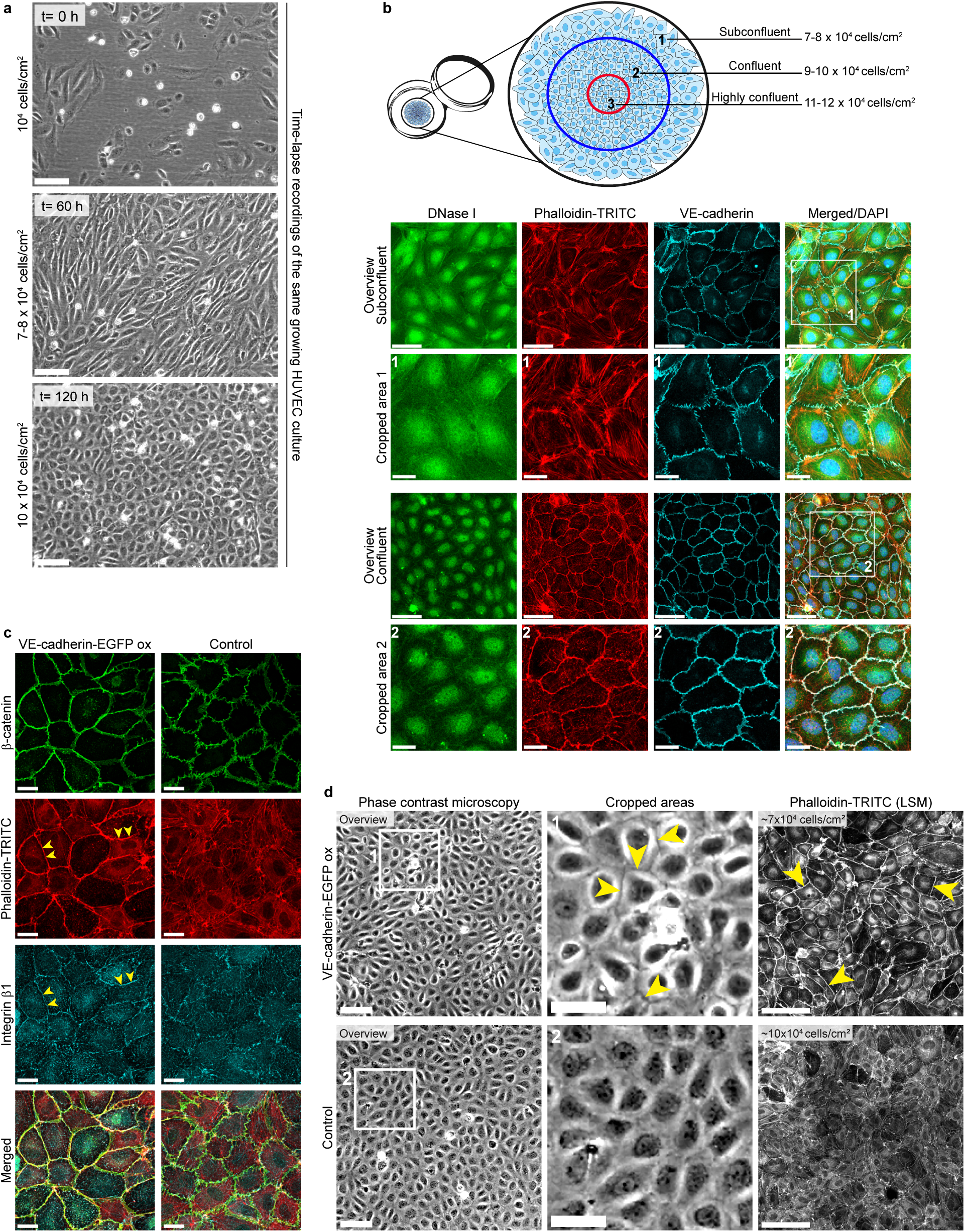
(**a**) The scheme illustrates the principle of the one-drop assay, which enables the analysis of different cell densities within the same cell culture to investigate G-actin levels. For this technique, a drop of cell suspension is placed at the center of a cross-linked gelatin-coated glass-bottom chamber. Due to the surface tension of the culture fluid, cells primarily assemble in the center, with decreasing cell numbers settling towards the periphery. Cultures displayed highly confluent (11-12 x 10^4^ cells/cm^2^), confluent (9-10 x 10^4^ cells/cm^2^), and sub-confluent (7-8 x 10^4^ cells/cm^2^) areas. Laser scanning microscopy (LSM) was used to examine the G-actin level with fluorescently labelled deoxyribonuclease I (DNase I) according to ^92^. VE-cadherin labelling served to identify cell borders, and filamentous actin was visualized with phalloidin-TRITC (overview: scale bar 50 µm; cropped areas: scale bar 20 µm). DNase I staining was performed as described. (**b**) Representative example of phase-contrast time-lapse recordings during cell growth to determine cell density-dependent dynamics. Images were taken at the same location in a growing HUVEC cell culture (scale bar: 100 µm). The cells were initially seeded at a density of 1 x 10^4^ cells/cm^2^. Images are shown at t = 0 hours, t = 60 hours, and t= 120 hours. The cell density at each respective time point is indicated. (**c**) LSM images of VE-cadherin-EGFP-overexpressing HUVECs and control cells, immunolabelled as indicated (scale bar: 20 µm). Arrowheads highlight prominent junctional actin and strong junctional integrin β1 recruitment in VE-cadherin-overexpressing cells. (**d**) Left panel (overview): Phase-contrast microscopy of VE-cadherin-EGFP-overexpressing HUVECs and control cells (scale bar: 100 µm). Middle panel: In the cropped area, arrowheads indicate enhanced contrast at cell borders in VE-cadherin-overexpressing HUVECs (scale bar: 50 µm), while controls did not show this phenomenon. Right panel: LSM overview of phalloidin-TRITC-labelled HUVECs, either with VE-cadherin overexpression or EGFP expression. Prominent junctional actin formation (arrowheads) is seen in VE-cadherin-EGFP-overexpressing cells, while stress fibers are largely absent (scale bar: 100 µm).

**Supplementary Figure 2:**
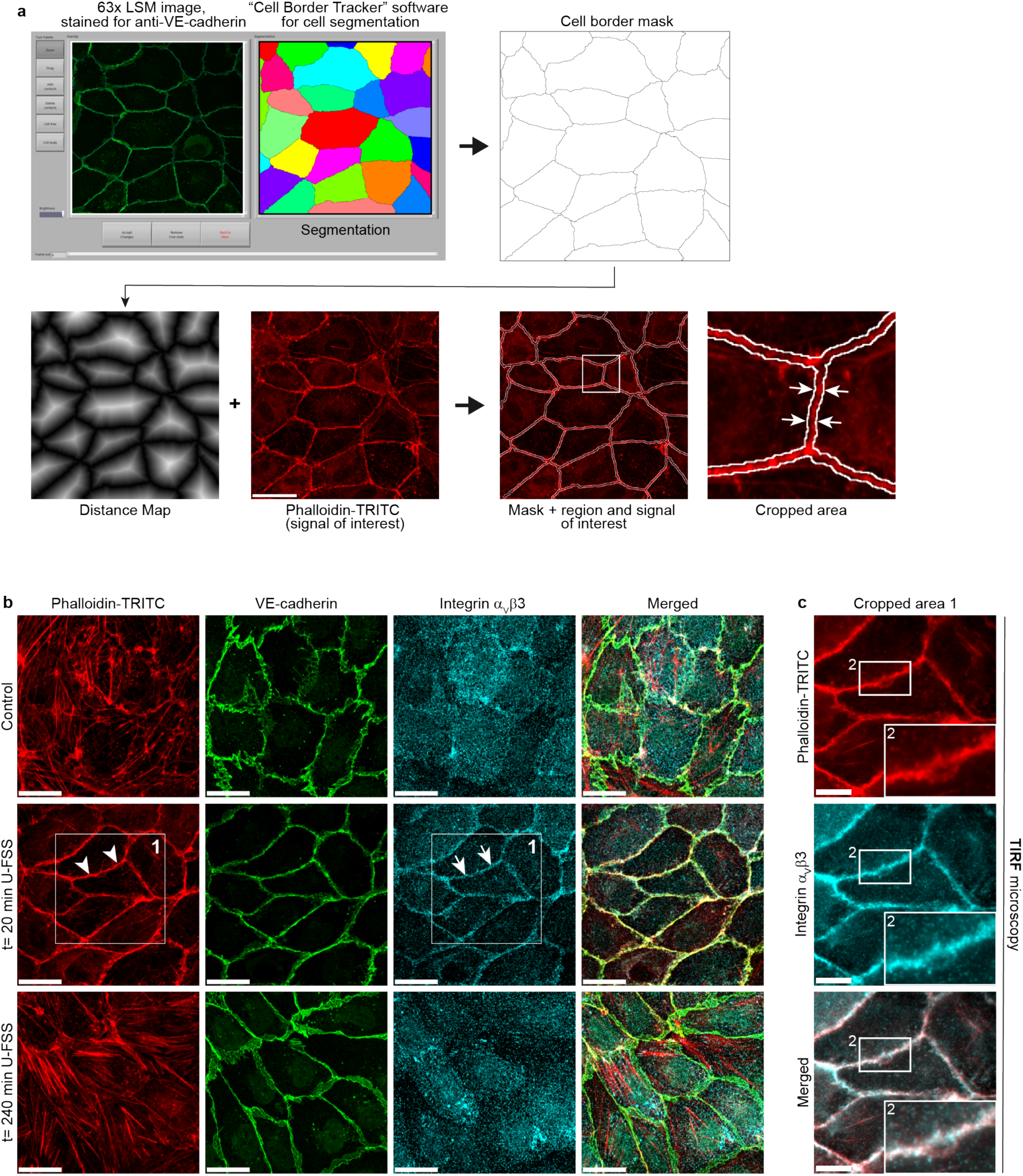
(**a**) Flowchart illustrating the measurement of fluorescence intensities at cell junctions using the CellBorderTracker (CBT) as described previously ^93^. CBT was used to acquire a cell border mask based on junction-localized anti-VE-cadherin staining. Next, a distance map was generated using Fiji ImageJ to define the region of interest (ROI), followed by relative actin intensity measurements. (Cropped area) The ROI was defined by extending 5 pixels on both sides of the segmentation line indicated by the white arrows. The relative junctional actin concentration was calculated as the integrated phalloidin-TRITC intensity within the ROI divided by the total cell border length of the ROI. (**b**) LSM images of confluent HUVECs subjected to 18 dyn/cm^2^ U-FSS for t = 0 min, t = 20 min, and t = 240 min, labelled for phalloidin-TRITC, VE-cadherin, and integrin α_V_β3 (scale bar: 20 µm). After t = 20 min of shear stress, a prominent junctional actin ring is formed, accompanied by integrin α_V_β3 recruitment. (**c**) Cropped areas 1 and 2 after t = 20 min of shear stress showing junctional actin and integrin α_V_β3 recruitment to the junctions, as verified by total internal reflection fluorescence microscopy (TIRF) (scale bar: 10 µm).

**Supplementary Figure 3:**
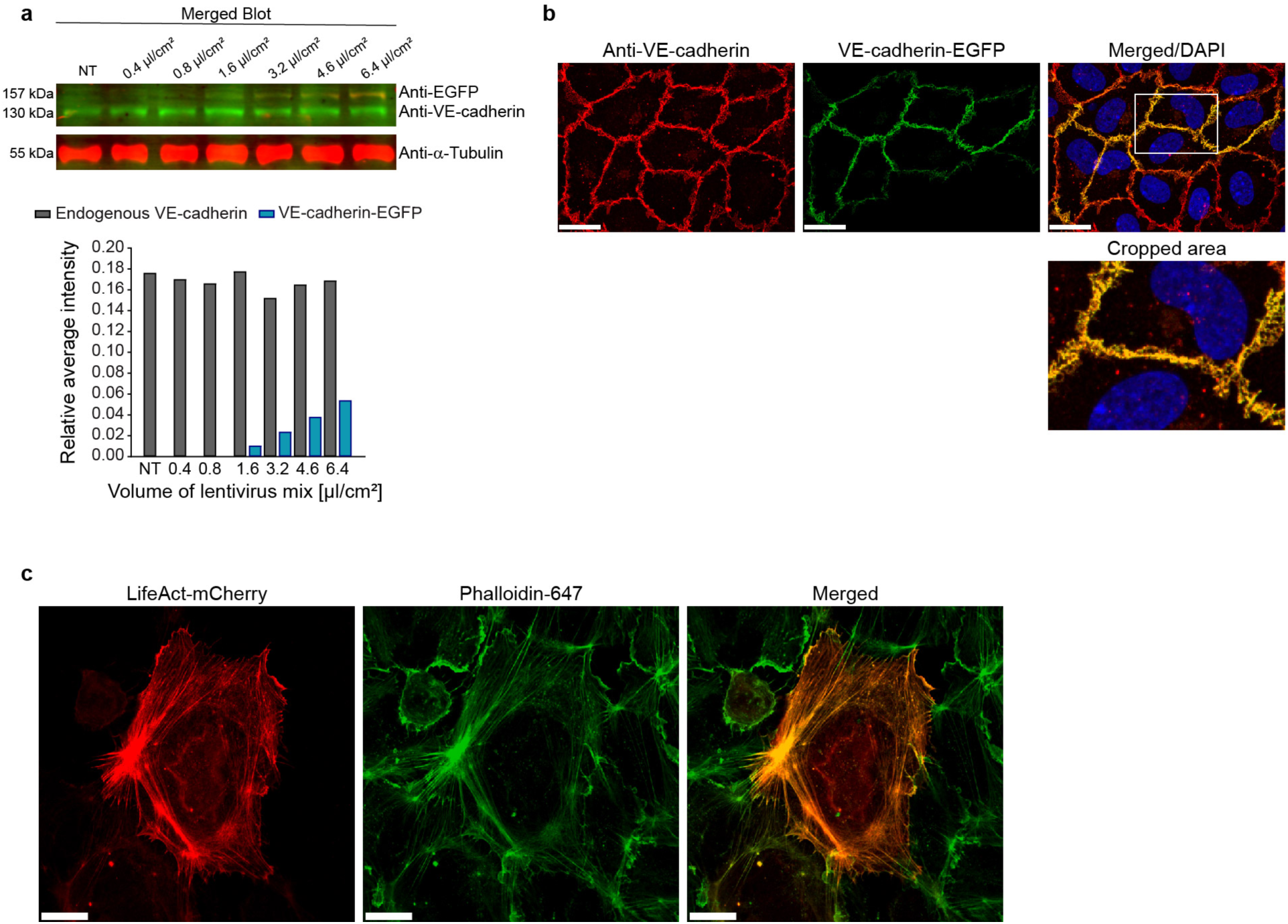
VE-cadherin-EGFP virus titration and LifeAct-mCherry expression. (**a**) Merged Western blot using anti-EGFP and anti-VE-cadherin antibodies. Samples were probed with different amounts of VE-cadherin-EGFP lentiviral mix for titration as indicated. α-Tubulin served as an internal loading control. (**b**) HUVECs expressing VE-cadherin-EGFP were stained with anti-VE-cadherin antibody and DAPI. VE-cadherin-EGFP colocalizes with the endogenous VE-cadherin pattern (cropped area). (**c**) HUVECs expressing LifeAct-mCherry were labelled with phalloidin-647.

**Supplementary Figure 4.**
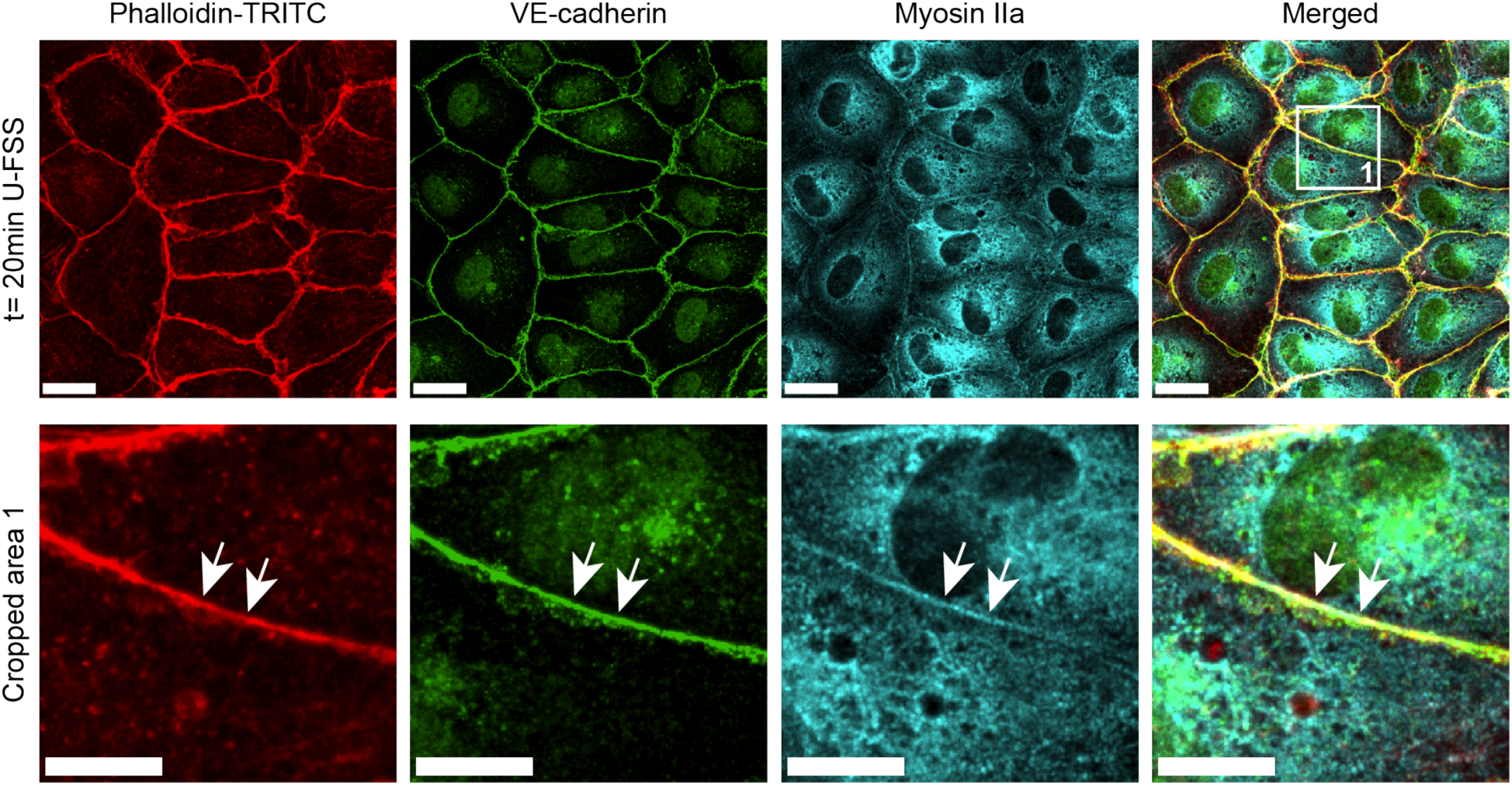
Laser scanning microscopy images of confluent HUVECs subjected to 18 dyn/cm^2^ U-FSS for t = 20 min and stained for filamentous actin (phalloidin-TRITC), VE-cadherin, and myosin IIa to demonstrate the presence of the contractile machinery at junctional VE-cadherin (scale bar: 20 µm). (Cropped area 1, scale bar: 10 µm) Arrows indicate the colocalization of myosin IIa and junctional actin.

**Supplementary Figure 5.**
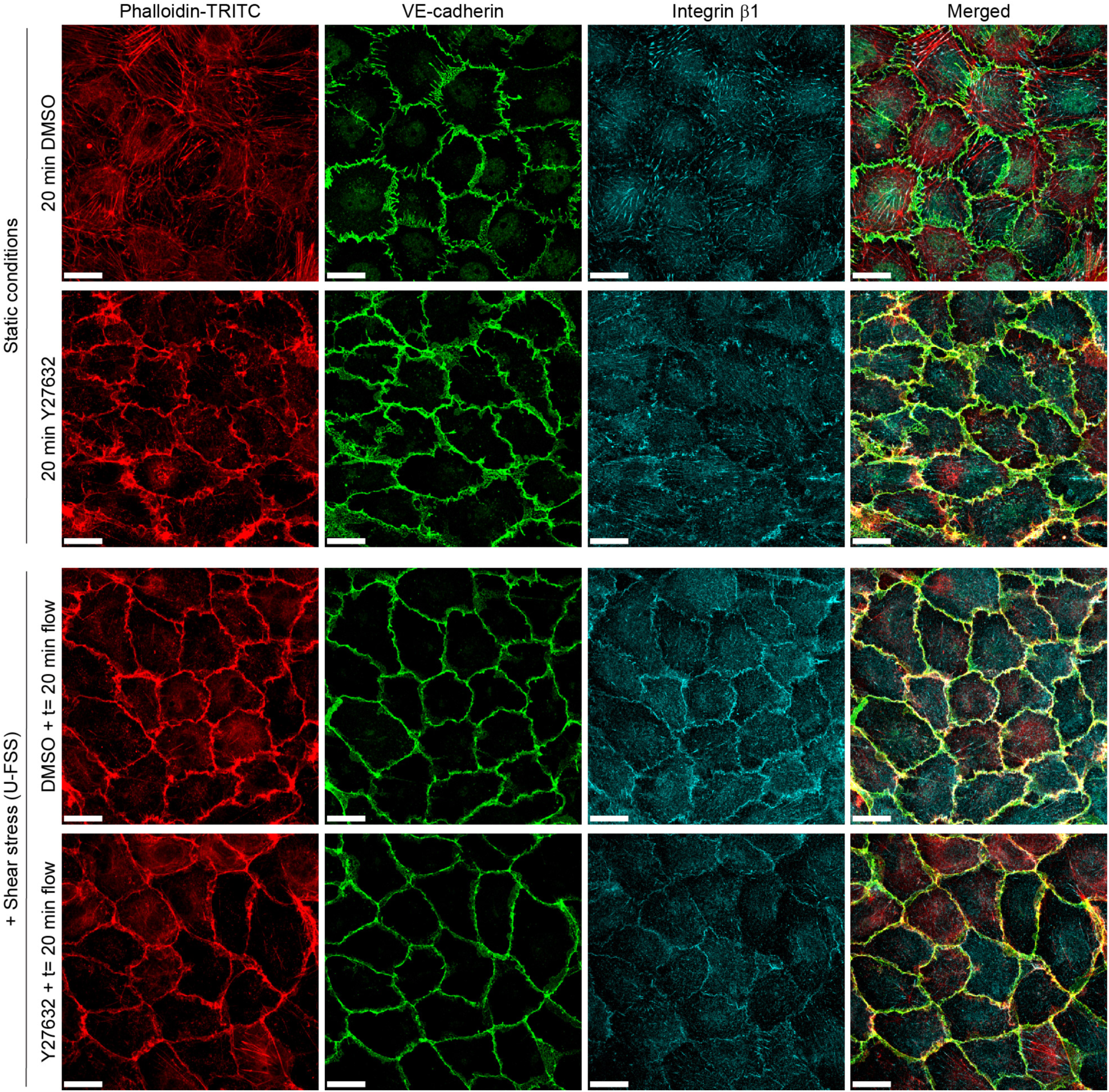
Laser scanning microscopy images of confluent HUVECs under static control conditions and after U-FSS (18 dyn/cm^2^) for 20 min, as well as after pre-incubation with Y27632 for 10 min combined with shear stress application, as indicated (scale bar: 20 µm). Cells were immunolabelled for actin filaments with phalloidin-TRITC, VE-cadherin, and integrin β1, as indicated. Y27632 treatment induced a heterogeneous VE-cadherin and actin distribution with both JAIL formation and a linearly organized VE-cadherin pattern associated with junctional actin formation. Cells pre-incubated with Y27632 and subsequently exposed to U-FSS for 20 min show a linear VE-cadherin pattern with prominent actin and integrin β1 overlap.

**Supplementary Figure 6.**
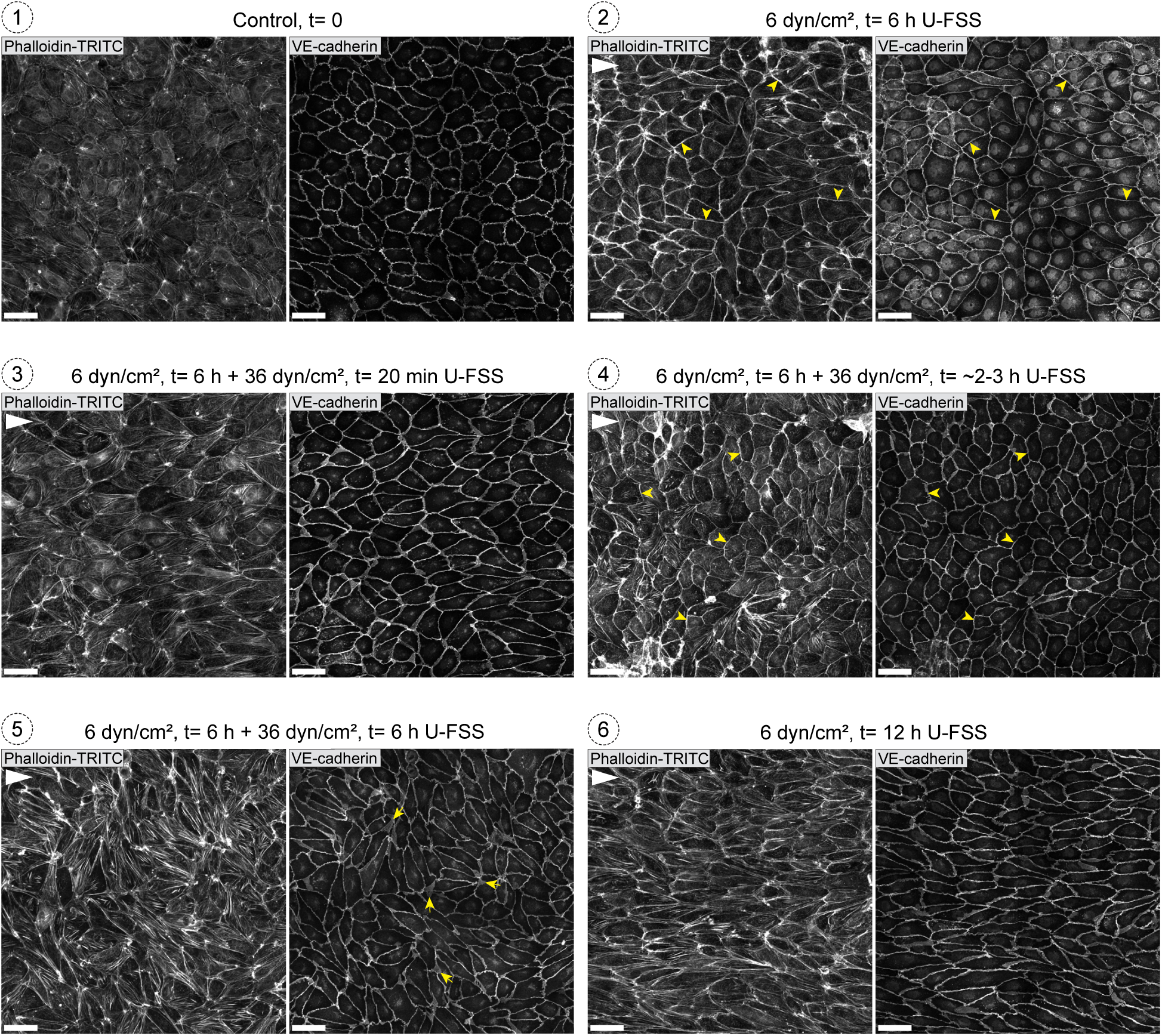
Laser scanning microscopy overview images (2×2 tiles, 40x magnification, 425 µm x 425 µm) of confluent HUVECs showing filamentous actin (phalloidin-TRITC) and VE-cadherin staining under different U-FSS profiles (scale bar: 50 µm). The white triangle indicates the direction of the U-FSS from the left. (**Image 1**) Control. (**Image 2**) HUVEC cultures after shear stress of 6 dyn/cm^2^ for 6 h. Note the linear VE-cadherin pattern (arrowheads) accompanied by a prominent junctional actin ring (arrowheads), while stress fibres are largely absent. (**Image 3**) After upregulation of the shear stress to 36 dyn/cm^2^ in shear stress-adapted cells, stress fibres re-emerge, accompanied by increased cell elongation. (**Image 4**) After approximately 2-3 h at 36 dyn/cm^2^, cells regain a more cobblestone-like morphology, characterized again by a linear VE-cadherin pattern and prominent junctional actin formation, which is associated with the almost complete disappearance of stress fibres. (**Image 5**) With continued shear stress at 36 dyn/cm^2^ U-FSS, cells develop actin stress fibres and align and elongate, driven by JAIL formation. (**Image 6**) Cells subjected to 6 dyn/cm^2^ for 12 hours also adapt over time, showing stress fibre formation and a heterogeneous VE-cadherin pattern, eventually leading to alignment and cell elongation.

